# Decoding Peripheral Tolerance: TCR Rules for pTreg differentiation in the Gut

**DOI:** 10.1101/2025.10.20.683415

**Authors:** Xinxin Chi, Charlotte H. Wang, Yollanda Franco Parisotto, William A. Nyberg, Vanja Cabric, Adelaide Gelineau, Yi Cao, David L Owen, Jonas Ambjörnsson, Diane Mathis, Justin Eyquem, Chrysothemis C. Brown, Christophe Benoist

## Abstract

Peripheral differentiation of regulatory T cells (pTregs) promoted by foreign antigens encountered in barrier tissues is considered a unique contributor to immunological tolerance to obligate non-self, like food or symbiotic microbes. The relative importance of adaptive recognition via the T cell receptor (TCR) vs environmental small-molecule or neuroimmune cues, is poorly understood. We leverage CRISPR-based TCR editing to perform in primary T cells *in vivo*, with a large panel of TCRs, a screen to assess pTreg differentiation induced by self, microbial, or dietary antigens. All antigen classes drive pTreg differentiation, which varies with the origin of the TCR: TCRs derived from Tregs enable pTreg differentiation much more effectively than those from Tconv. TCRs recognizing self, microbial, or dietary antigens elicit distinct pTreg phenotypes, Helios⁺, RORγ⁺, or both. Mechanistically, these trace to different types of antigen-presenting-cell involved. That Treg-derived TCRs preferentially drive tolerogenic fate speaks to preferential drivers of tolerogenic therapy.

---

The gastrointestinal tract represents a unique and complex immunological environment, constantly interacting with a diverse array of dietary antigens, commensal microbes, and self-antigens. Maintaining immune homeostasis in this dynamic setting relies on appropriate immune responses to microbes and food antigens, in the context of their broad communities^1,2^. CD4⁺ T cells play a central role in these processes, and different microbes have been shown to elicit distinct responses ^3–8^. Among those, regulatory T cells that express the transcription factor FoxP3 (Tregs) are essential for suppressing excessive inflammation and fostering tolerance in the gut^1,9^.

Many Treg cells differentiate in the thymus (tTregs), driven by recognition of self-antigens, but another pool differentiates in peripheral lymphoid organs (pTregs) from conventional CD4⁺ T cells (Tconv), a conversion driven by foreign antigens, in particular from food and microbes in the gut ^9,10^. These two populations have fundamentally different operational logics, since one is focused on *self* antigens initially encountered in the thymus, while the other mostly deals with *foreign* antigens. Accordingly, tTreg and pTregs have been shown to play complementary parts in immunoregulation ^11–14^. Understanding what drives T cells towards effector or pTreg differentiation is key to elucidating peripheral immune tolerance and to manipulate T cell responses for cell therapy and vaccination.

In gut-associated lymphoid tissues (GALT), the different flavors of Treg cells can schematically be categorized into Helios⁺ (also Gata3⁺) and RORγ⁺ (also cMaf⁺) categories of Tregs, as well as Helios⁻RORγ⁻ double-negatives (DN). RORγ⁺ Tregs are strongly tuned by gut bacteria ^6,15–18^. They control intestinal inflammation, IgA production, and food allergies ^6,13,15,19,20^, and are only partially dependent on FoxP3 ^13,21^. In contrast, Helios⁺ Tregs strictly require FoxP3, are thought to partake in tissue repair via their production of amphiregulin ^22–25^ but also curtail inflammation more generally. Dietary proteins have been reported to regulate DN Tregs in small intestine ^26^. It was originally thought that Helios⁺ Tregs are tTregs that differentiate in the thymus, while RORγ⁺ Tregs are pTregs that differentiate from Tconv cells in the periphery. Transfer and lineage tracing experiments in several systems supported this notion ^13,17,27,28^. On the other hand, some pTregs can be Helios⁺ ^28^, and analyses of αβTCR sequencing data revealed clonotypes shared by RORγ⁺ and Helios⁺ Tregs ^28,29^, which implies that they share some common origin. These are not compatible with a strict Helios⁺ = tTreg, RORγ⁺ = pTreg, equivalency, and this important question remains unresolved.

The functional importance of pTregs in balancing effector T cell responses to the same foreign antigens is well understood, but it is unclear what determines the conversion of a Tconv cell into a pTreg ^9^. Are the effects of microbiota and diet on gut Treg cells direct (antigen-dependent) or indirect (via metabolites and other cells)? The question is made more complex by the diversity of actors that influence colonic Treg cells: microbes have a strong influence, as do antigen presenting cells (APC) and in particular the recently described RORγ⁺ APCs ^30–32^. Neuroimmune connections are involved, and we showed that Trpv1⁺ sensory neurons modulate RORγ⁺ Treg cells ^7,33,34^. The characteristics of the antigens that can drive pTreg generation are also unclear: is it simply a matter of a cell being activated by a strong agonist in peripheral tissues? If so, any Tconv cell should have the ability to become a pTreg. Another model posits that more specific cues, encoded in its TCR, determine a T cell’s ability to become a pTreg – for instance by signaling with a particular intensity range, or by recognizing antigen presented by a particular type of antigen-presenting cell.

These questions have been difficult to address because of the limited range of TCRs that can be explored with transgenic models. Individual transgenic models support pTreg generation with variable efficacy ^17,35^, but it is unclear how to generalize from one-off observations. Recent work from our lab analyzed in detail the phenotypes and TCRs of colonic CD4⁺ T cells in mice colonized by single microbes, identifying thousands of individual clonotypes ^29^. Some of these were shared between colonic Treg and Tconv cells, suggesting involvement in pTreg cells. Here, by leveraging this data store and a powerful system for efficient TCR engineering in primary mouse T cells, we performed a mini-screen to assess the ability of a diverse panel of TCRs to drive pTreg differentiation. The results reveal that all TCRs are not equal. The ability to become a pTreg, and to adopt the RORγ⁺ phenotype, are both deeply pre-determined by a cell’s TCR, by the class of antigen that it recognizes, and by the type of APC which presents it.

## RESULTS

### Colonic CD4 T cell TCRs are activated by food, bacterial, and self antigens

To understand the potential of TCRs from colonic CD4 in shaping cell destiny, we selected from our previous dataset ^29^ a panel of 24 TCRs derived CD4⁺ T cells from colonic lamina propria of gnotobiotic mice monocolonized with *C. ramosum* (Crm) *or E. coli Nissle* (EcN). These TCRs originated from several cell-types (Tconv, Treg of different phenotypes (RORγ+, Helios⁻, or RORγ⁻Helios⁻ (DN)) (**Fig. S1**). To functionally express these TCRs in normal T cells, we utilized a CRISPR-Cas9-based TCR knock-in system recently established by some of us^36^. This system utilizes the AAV variant Ark313, evolved to infect primary mouse T cells. It delivers to cells from Cas9-expressing transgenic mice, in a single vector, gRNAs that inactivate endogenous *Trac* and *Trbc* loci and a template for homology-directed repair targeting *Trac* that inserts the sequences coding for the TCR of interest under the control of the endogenous promoter (as a 2A-spaced bicistronic construct; **Fig. 1A**). The endogenous *Trac* and *Trbc* genes are inactivated in ∼90% of transduced primary CD4⁺ T cells, 20-40% of which express the desired TCR, detectable by the appropriate TCR-V-specific monoclonal antibodies (illustrated for two such constructs in **Fig. 1B**; note that the process leaves in the cultures a few T cells non-edited cells, as well as TCR-negative cells, which provided useful internal standards in the experiments below).

**Figure 1.**
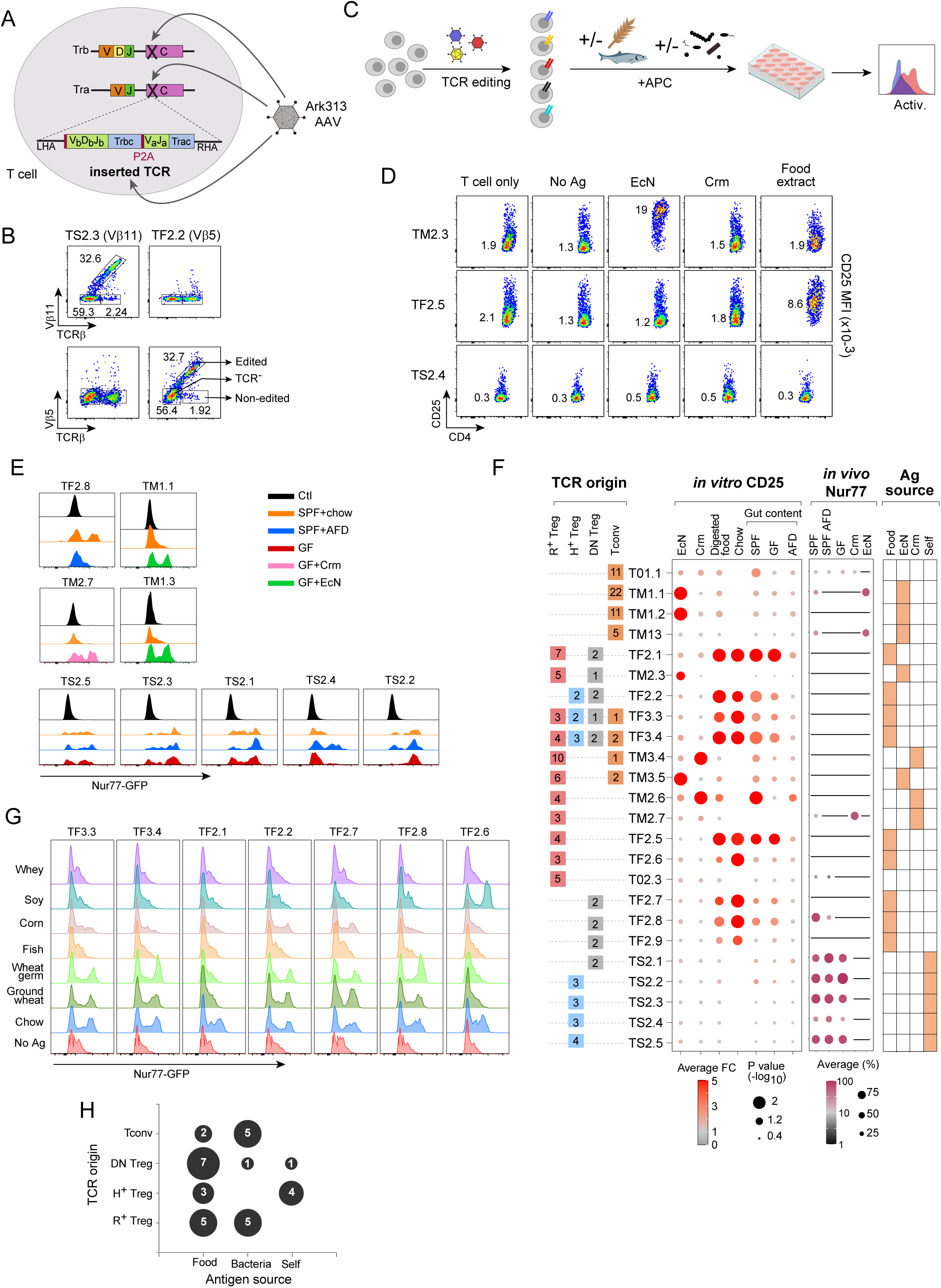
Reactivity of TCRs derived from colonic Treg and Tconv cells. **(A)** Diagram illustrating TCR editing with the Ark313 AAV vector. The AAV encodes 2 sgRNAs to cut into the C regions of TCRα and TCRβ loci, as well as an expression cassette for the TCRα and TCRβ of interest (connected by a 2A linker) that inserts by homologous recombination into the cleaved TCRα region **(B)** Representative flow cytometric plots showing TCRVβ11 and TCRVβ5 expression in CD4⁺ T cells edited to express TS2.3 (Vβ11⁺) or TF2.2 (Vβ5⁺) TCRs. **(C)** Schematic experimental design: CD4⁺ T cells are edited to express different TCRs, and cultured with APCs supplemented with different sources of intestinal antigens, before readout of activation markers by flow cytometry. **(D)** Flow cytometric plots of CD25 expression in TCR edited cells co-cultured with the indicated antigens, representative of 2 or more experiments. Numbers are Mean Fluorescence Intensity (MFI) **(E)** Representative experiment: CD4⁺ T cells from Nur77-GFP/Cas9 mice were TCR-edited and transferred into the indicated hosts. After 24 hrs, mLN (gated on donor-derived Edited cells) were analyzed for activation denoted by GFP expression. “Ctl” controls were TCR-negative cells from those transfers. **(F)** Summary of the characteristics and reactivity of all TCRs used in this study. From left to right: summary of the type of colonic CD4⁺ T cell in which the TCRs were originally discovered^29^, with numbers of cells expressing these clonotypes; compilation of the *in vitro* CD25 induction experiments, dot plot displaying average fold change (FC) and –log10 (p-value) of CD25 MFI relative to APC-only controls; compilation of in vivo Nur77-GFP⁺ induction experiments, as % GFP⁺ cells among TCR-edited cells – black line: not done; a summary of the antigen specificity for all TCRs used in this study. **(G)** TCR-edited Nur77-GFP/Cas9 CD4⁺ were co-cultured with APC and protein extracts for 6 hours; plots of GFP expression are representative of 2 or more experiments. **(H)** Summary plot connecting the TCRs’ antigen specificity with their cell-type of origin (dots sized according to the number of instances – clonotypes found in several cell-types counted more than once).

To assess the antigen specificity of the TCR panel, we co-cultured TCR-edited CD4⁺ splenocytes with APCs loaded with different complex antigens: heat-killed bacteria, protein extract from colon content, solubilized mouse chow, intestinal content (from SPF or germfree mice, fed standard chow or a protein-antigen-free (amino acid based diet, hereafter AFD) (**Fig. 1C**). CD25 induction was used as an indicator of responses **(Fig. 1D, Table S1**). Specific reactivity to microbial (EcN or Crm) or food antigens was readily identified in this manner for 15 of the 24 TCRs tested, including TCRs emanating from Tconv or various Treg subsets (**Fig. 1D**, tabulated in **Fig. 1F**), indicating that colonic CD4⁺ T cells include a very high frequency of T cells reactive to gut luminal antigens. In contrast to a previous report^37^, we did not detect polyreactive T cells that respond to antigens from several microbes, but this may result from different microbiota (community vs monocolonized) in mice from which the TCRs emanated.

This simple *in vitro* activation system might miss reactivities to some antigens that would require processing and presentation by APCs *in vivo*. We set an *in vivo* test system, wherein the TCRs whose antigen specificity could not be determined *in vitro* were edited into T cells from Nur77-GFP reporter mice, in which GFP expression denotes TCR stimulation ^38,39^. Confirming *in vitro* results, cells expressing the food-reactive TF2.8 TCR upregulated Nur77-GFP expression in normal but not AFD-fed hosts, and cells expressing the TM1.1 TCR were activated in EcN-colonized but not in SPF hosts (**Fig. 1E, F**), In this assay, five TCRs (TS2.1-TS2.5) that seemed unresponsive *in vitro* proved to be activated *in vivo* in all hosts, irrespective of microbe (GF) or food antigens (AFD), indicating responsiveness to self-antigens rather than bacterial or dietary antigens (**Fig. 1E, F, Table S1**). No antigen reactivity could be identified for T02.3 and T01.1 TCRs. Thus, by combining *in vitro* co-culture and *in vivo* assays, we identified 8 microbe-reactive, 9 food-reactive and 5 self-reactive TCRs (**Fig. 1F**).

Although we did not aim here to formally identify the actual peptides recognized by these food-reactive TCRs, it was of interest to assess whether they corresponded to a single protein type or immunodominant epitope. Hence, we cultured TCR-edited Nur77-GFP reporter cells with APCs and extracts from specific components of normal mouse chow, which identified several different patterns: reactivity to soy extract for one TCR, to wheat for five other TCRs, of which two cross-reacted with corn (**Fig. 1G**). Thus, the TCRs in our panel that react to dietary antigens recognize a variety of epitopes, although dominated by grain proteins, a diversity comparable to that recently reported^40^.

Integrating these response data showed that the types of colon T cells from which these TCRs originate was clearly related to their specificity (**Fig. 1F**) as further summarized in **Fig. 1H**. Self-reactive TCRs mostly originated only from Helios⁺ Tregs. Microbe-reactive TCRs stemmed from either Tconv or RORγ⁺ Tregs, consistent with the types of cells known to be microbe-reactive ^6,20,41–44^. Food reactive TCRs derived from the broadest array of colonic cells, Tconv and all Treg subsets, arguing for a broad phenotypic dispersion of the immune response to food antigens.

### TCR-antigen recognition directs T cell fate and tissue distribution

Having decoded the categories of antigens that activated the panel of colon TCRs, we then asked how TCRs reactive to different antigens would drive the differentiation of CD4⁺ Tconv cells (clonal proliferation, pTreg generation?). Congenically-marked (Thy1.1) spleen Tconv cells were edited as above to express individual TCRs from our panel, their FoxP3-negative status verified (**Fig. S2A**), and transferred into normal host mice in which the cognate antigens were present or absent (**Fig. 2A**; for food-specific TCRs, mice fed regular chow vs AFD diet; for microbe-specific TCRs, SPF or GF mice colonized or not with EcN or Crm). To streamline this screen, cells expressing different TCRs were often multiplexed for transfer into the same host (**Fig. S2B**), which also enhanced the comparability between different TCRs. Thy1.1⁺ donor cells were identifiable ten days later in different locations, in systemic lymphoid organs (spleen, subcutaneous lymph nodes – scLN) and in the GALT (small intestine– and colon-draining mesenteric LNs – smLN and cmLN, respectively –, colon and ileum lamina propria (**Fig. S2C**). For quantitation, we normalized their numbers to those of the co-transferred TCR-negative cells (and relative to this ratio at the time of transfer) to reflect cognate TCR-driven cell accumulation in each tissue (**Fig. 2B**). Representative results for one TCR are shown in **Fig. 2C**, but several important findings emerged from the fully integrated results (23 TCRs in 155 independent transfers, **Fig. 2D** **and Table S2**). First, and irrespective of specificity (food, microbe or self) virtually all of these TCRs drove preferential accumulation in GALT locations, and a relative depletion in systemic lymphoid organs. This accumulation was antigen-dependent, largely disappearing in Ag-free control hosts. The exceptions were T01.1, which induced a run-away expansion in all tissues, and TM3.5, which showed comparable enrichment in all tissues. Second, different TCRs led to preferential distribution in the intestinal tissues themselves (colon, ileum, TF2.2, TS2.2) or in the draining LNs (TM2.6, TF2.6; **Fig. 2D**). Moreover, individual TCRs displayed organ-preferential patterns. For example, TF2.1-expressing cells preferentially accumulated in the ileum and its draining smLN, whereas TS2.4-expressing cells were more enriched in the colon and its draining cmLN.

**Figure 2.**
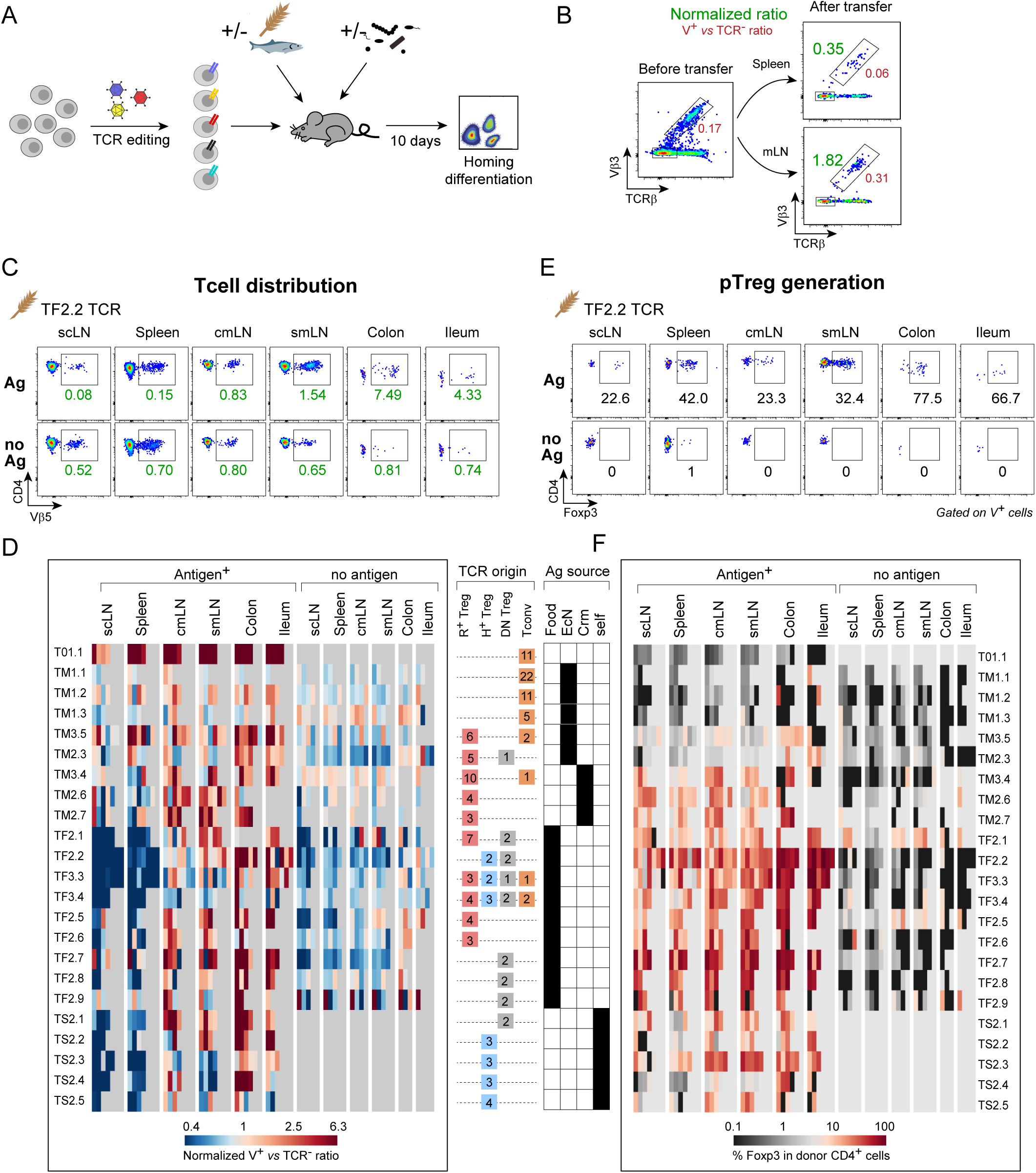
TCR-antigen recognition directs T cell tissue distribution and pTreg generation. **(A)** Diagram of experimental design: CD4⁺ Tconv cells were TCR-edited as above, and transferred into host mice which provided, or not, the corresponding antigen sources. For food antigens: SPF mice fed regular chow or AFD diet; for microbial antigens SPF mice gavaged or not with *E. coli Nissle*, or germfree mice gavaged, or not, with or without *C. ramosum*; for self-reactive TCRs, SPF mice – no controls possible). Donor-derived cells in different tissues were analyzed 10 days after transfer. **(B)** Representative plots depicting the normalization method used. After editing to express the TF2.8 TCR (Vβ3⁺), the donor cell population was normalized to TCR-negative cells in each pool (red numbers; note how it increased in the mLN relative to spleen), and their relative abundance in each organ was further normalized to before-transfer values (green numbers), in order to enable comparison across different experiments. **(C)** Representative flow cytometric plots of tissue distribution of TF2.2-expressing cells in different organs, 10 days after transfer (scLN: subcutaneous LN; cmLN: colon-draining LN; smLN: SI-draining LN) green numbers: normalized representation in each tissue, calculated as detailed in B. **(D)** Compilation of tissue distribution of all transfer experiments as in C, representing the normalized ratio of TCR-edited cells (V⁺) vs TCR⁻ cells, across different tissues. Results in recipients devoid of antigen (A) are shown at right. Each bar an individual host, n= 3-6, with a few dropouts because cell numbers were too low for quantitation. A recall from Fig. 1 of TCR origin and specificity is shown at right. **(E)** Representative flow cytometric plots of FoxP3 expression in TF2.2-expressing cells, 10 days after transfer. **(F)** Compilation of FoxP3 expression in donor cells (as in E) for all transfer experiments, in antigen-positive or negative hosts, color-coded as the percent FoxP3⁺ cells among Edited⁺CD4⁺ T cells. Each bar represents an individual mouse (n= 3-6). Only samples with more than 10 cells for each TCR-edited population were included in the analysis shown in D and F.

We then examined the capacity of individual TCR-antigen pairs to direct pTreg differentiation in the same transfers. As illustrated in **Fig. 2E** for the TF2.2 TCR, FoxP3⁺ pTregs were detected abundantly ten days after transfer, not only in the GALT but also in systemic lymphoid organs. pTreg differentiation was completely abrogated in the absence of antigen (**Fig. 2E**, bottom row). The full data (**Fig. 2F, Table S2**) confirmed this dependency. Furthermore, essentially none of the co-transferred TCR-negative or non-edited cells developed FoxP3 expression (**Fig. S2D**), showing that a TCR-dependent trigger was required. There was, however, substantial variation in the ability of different TCRs to support pTreg differentiation, as illustrated in **Fig. 2F**, which reveals more several trends. First, TCRs originally identified exclusively in Tconv cells (T01.1, TM1.1, TM1.2, TM1.3, top rows in **Fig. 2F**) generally failed to induce Tregs, no matter in monocolonized gnotobiotic mice or SPF mice gavaged with EcN following streptomycin treatment (TM1.1, TM1.2 in **Fig. S2E**), while all those originally from Tregs did so, if to varying extents. Second, TCRs that recognize food antigens generally promoted pTreg more robustly than those reactive against microbial or self-antigens —a trend observed in both EcN monocolonized mice and SPF mice **(Fig. S2E**). Third, more pTreg differentiation was observed with TCRs reactive against Crm than against EcN, in keeping with Crm being one of the strongest Treg inducer in our previous monocolonization studies ^6^.

TCRs originally derived from colonic Tregs reactive against self-antigens drove pTreg differentiation with a preferential tissue localization in the GALT (**Fig. 2F**), which seemed paradoxical given that self-reactive Treg cells are usually thought to originate in the thymus. To rule out reactivity against a food or microbial antigen that would have escaped our earlier assays, we transferred edited Tconv cells into hosts devoid of microbial (germfree), or dietary (AFD), or both (AFD-fed germfree) sources of foreign antigens. Except for TS2.5, pTreg generation was comparable in all hosts, confirming that true self-reactivity is involved (**Fig. S2F**). These results suggest that reactivity against self-antigens, apparently specific of the gut but unrelated to dietary or microbial antigens, is able to drive pTreg differentiation.

pTregs are generally considered to be specific to mucosal tissues, and it was surprising to find them disseminated in so many locations. Although lower in proportion, pTregs in the spleen do form large contingents (roughly 25% of the pTregs generated in any one mouse are found in the spleen (**Fig. 2E, F**)). To rule out that these splenic pTregs might be due to the particular conditions of our Cas9/AAV editing system, we performed parallel transfers with Tconv cells from two lines of transgenic mice (BθOM and HH7-2) that express TCRs reactive against antigens from *Bacteroides thetaiotamicron* and *Helicobacter hepaticus*, respectively ^8,17^. As with the transfer of TCR-edited cells above, pTregs from these transgenically-encoded TCRs were detected in various tissues, including the spleen and scLNs, but only in microbe-carrying hosts (**Fig. S2G**), ruling out a quirk of the TCR editing system. HH7-2tg TCR transgenic cells exhibited higher pTreg generation than BθOM cells, reflecting again the differential ability of TCR/antigen/APC combinations to support pTreg generation.

To determine where pTregs first appear, we traced their generation over time. For food– and self-antigen-reactive TCRs, pTreg formation was first observed in the mLN as early as two days post-transfer, conversion of cells expressing microbe-reactive TCRs exhibiting a slower response (**Fig. S3A-B**). pTregs only accumulated a little later in the spleen. Together with the data above, we interpret these kinetics to mean that differentiation first occurs in the mLN, with secondary spread to the spleen or to intestinal tissues, a balance controlled in part by availability of antigen in different locales.

Collectively, these findings suggest that pTreg can be induced by food, bacteria or self-antigens, widely distribute in hosts and are not restricted to gut-associated tissues.

### pTreg phenotypes are determined by the TCR

As introduced above, colonic Treg cells exist in functionally different phenotypes (RORγ⁺, Helios⁺ or DN), and the relationship between these phenotypes and Treg origin (pTreg and tTreg) is debated. We thus leveraged our panel of TCRs to assess the phenotypes adopted by pTregs elicited by individual TCR-antigen pairs after transfer of edited cells into antigen-carrying hosts. Several important observations stood out. First, pTreg cells generated by antigen-driven conversion from transferred Tconv cells can adopt several different phenotypes: predominantly Helios⁺ for TS2.3 (albeit with slightly lower levels of Helios expression than in host’s Treg cells), predominantly RORγ⁺ for TM2.7, both for TF2.2 (**Fig. 3A, S4**). All pTregs also included a strong contingent of DNs. These results clearly established that pTregs are not obligatorily RORγ⁺. Secondly, these phenotypes correlated strongly with the class of antigen recognized by the TCRs. Microbes predominantly induced RORγ⁺ pTregs in the mLN, as also visible in colon and spleen (**Fig. 3B**), while self-reactive pTreg exclusively expressed Helios, with no RORγ expression detected in any tissue. In contrast, dietary antigens induced pTregs exhibit a mixed phenotype, expressing either RORγ or Helios in the mLNs or spleen, with a shift to higher RORγ expression pattern in colon (**Fig. 3A-B**). These results, integrated in **Fig. 3C**, indicate that pTreg phenotype is shaped by the TCR-antigen pair, and suggest further molding by the colon microenvironment. Thirdly, as explicitly shown in **Fig. 3D**, the phenotype adopted by the pTregs was also strongly conditioned by the phenotype of the cell from which we originally isolated the αβTCR: RORγ⁺ pTregs were mostly induced by TCRs originally isolated from RORγ⁺ pTregs. This link again supports the notion that the TCR encodes, likely by the antigen specificity, the nature of the APCs that present this antigen, and the environment in which activation occurs, the phenotype of the resulting pTreg.

**Figure 3.**
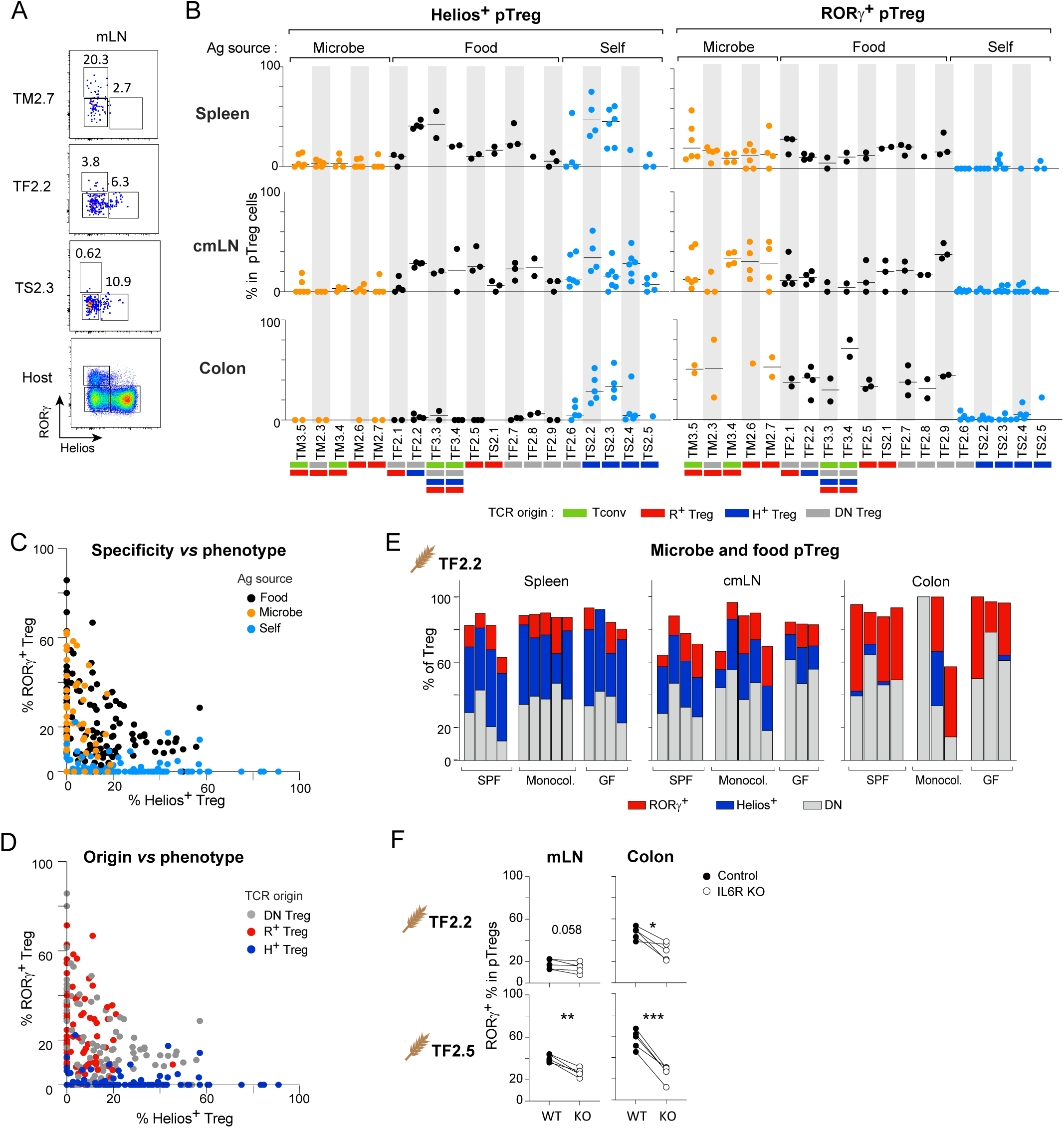
pTregs can adopt either RORγ vs Helios phenotypes, driven by TCR specificity and origin. **(A)** Representative plots showing RORγ vs Helios expression patterns in TCR-edited pTreg cells in mLN (day10, transfers as in Fig. 2), gated on CD4⁺Edited⁺FoxP3⁺ cells. **(B)** Compilation of Helios⁺ or RORγ⁺ frequencies (as a proportion of pTregs) in spleen, colonic mLN and colon, grouped according to TCR specificity (from the same experiments as Fig. 2C-F, S2E-F). **(C)** Aggregated data (mean of % Helios⁺ vs mean or RORγ⁺ among pTregs), color-coded according to antigen specificity. **(D)** As C, color-coded for the TCR cell of origin. **(E)** Proportions of RORγ⁺, Helios⁺ and RORγ⁻Helios⁻ (DN) cells among TF2.2 pTregs in SPF, gnotobiotic mice monocolonized with EcN or Crm, or germ-free hosts. **(F)** The *Il6ra* locus in CD4⁺ T cells (congenically tagged with Thy1.1 or Thy1.1/2) was targeted by electroporation of Cas9/gRNA complexes^46^ (or irrelevant gRNA as control), followed by standard TCR editing procedure before transfer into CD45.1/B6 hosts for 10 days. RORγ⁺ percent among pTregs were quantified in control or IL-6-receptor deleted cells. *P < 0.05, **P < 0.01, ***P < 0.001 by paired Student’s t test. Each symbol or bar in (B) to (F) represents an individual mouse. Only samples with more than 5 pTregs per TCR-edited population were included for analysis.

In normal mice, RORγ⁺ Tregs are highly represented in the colonic LP, relative to mLN and other tissues, and their proportions are modulated by microbes, as shown by gnotobiotic and antibiotic experiments ^6,15^. We thus asked whether microbes would alter the proportion of RORγ⁺ pTregs, even for cells whose differentiation was driven by a food antigen. We compared RORγ⁺ frequencies among pTregs expressing the food-reactive TF2.2 TCR, after transfer in hosts carrying different microbiota: normal complex microbiota (SPF), single microbe (monocolonized with EcN or Crm), or none (germ-free; **Fig. 3E**). The distribution of pTreg phenotypes, high RORγ⁺ in colon, high Helios⁺ in spleen remained consistent in these conditions (**Fig. 3E**). Thus, food antigens can drive the differentiation of RORγ⁺ pTregs, without requiring contributions from gut microbes.

Several studies from our group and others have found that IL-6 ^6,15,28,33^, including IL-6 derived from neurons ^45^, plays an important role in tuning the population of RORγ⁺ Tregs in the colon, but it was unclear whether this reflected an influence on pTreg differentiation or on a homeostatic setpoint. To address this question, we implemented a sequential editing strategy of primary Tconv cells, first inactivating the *Il6ra* locus by electroporation of Cas9/gRNA complexes ^46^, prior to TCR editing as above. We then transferred these cells, as an internally-controlled co-transfer of TF2.2 or TF2.5-edited Tconv, also edited with control or *Il6ra*-inactivating gRNAs, into the same hosts. The results showed that inactivating the IL-6 receptor in Tconv cells did not affect pTreg differentiation overall, but significantly reduced the fraction of RORγ⁺ pTregs (**Fig. 3F**). Thus, IL-6 signaling plays a crucial role in promoting RORγ expression in food-specific pTregs.

### Different antigens induce transcriptionally unique pTregs

These results showed that different classes of gut antigens led to different pTreg generation, numerically and phenotypically. To understand more fully if these different classes also led to phenotypically distinct Treg populations, we performed single cell RNA-sequencing on newly generated pTreg cells expressing TF2.2 (food-), TM3.5 (microbe-) or TS2.3 (self-reactive) TCRs, 10 days after transfer (sorted based on FoxP3-GFP expression from pooled mLNs). We also analyzed those edited and transferred cells that did not convert to FoxP3-positivity, as they might inform on what was missing in cells that did not become pTregs, and/or on transition intermediates. Non-edited and TCR-negative donor cells, which went through the same culture and transfer steps but received no differentiation-inducing signals, were used as negative controls. As references for normal Treg and Tconv cells, we included CD25hi Tregs from the host mice as well as GFP⁺ and GFP⁻ cells from an age-matched *FoxP3-gfp* mouse (all multiplexed by hashtagging into the same 10X Genomics run). The overall distribution of all cells in this experiment is shown in the Uniform Manifold Approximation and Projection (UMAP) of **Fig. 4A** (data from a very similar replicate experiment in Fig. S5). Tconv, rTreg and aTreg clusters were distinguished by the usual markers (**Fig. 4A-B** and **S5A-B**), normal gut Treg and Tconv cells from the host defining much of this space, reproducible in different mice (**Fig. 4C** **and S5C, bottom row**), as did donor-derived non-edited and TCR-negative cells. Strikingly, donor-derived pTreg cells mapped to specific locations, slightly different for each of the three TCRs (**Fig. 4C** **and S5C,** better compared in **Fig. 4D** **and S5D**). Self-reactive TS2.3 pTregs formed a tight cluster among aTregs. Microbe-reactive TM3.5 pTregs were more dispersed in the Treg space, while food-reactive TF2.2 pTregs included a main aTreg cluster close but distinct from TS2.3 pTregs as well as cells in the upper RORγ⁺ region. These UMAP differences also corresponded to variations in the expression of typical Treg marker genes like *Il2ra*, *Ctla4 or Tnfrsf4* (**Fig. 4E** **and S5E**). Moreover, the RORγ⁺ Treg signature ^47^ was enriched in TF2.2 pTregs compared with TS2.3 pTregs (**Fig. 4F** and **TableS3**). Thus, pTreg differentiation leads to marked aTreg phenotype (consistent with recent activation by antigen), among which the TCR/antigen combinations lead to distinct outcomes. The same dispositions were observed in the replicate experiment, **Fig. S5C**).

**Figure 4.**
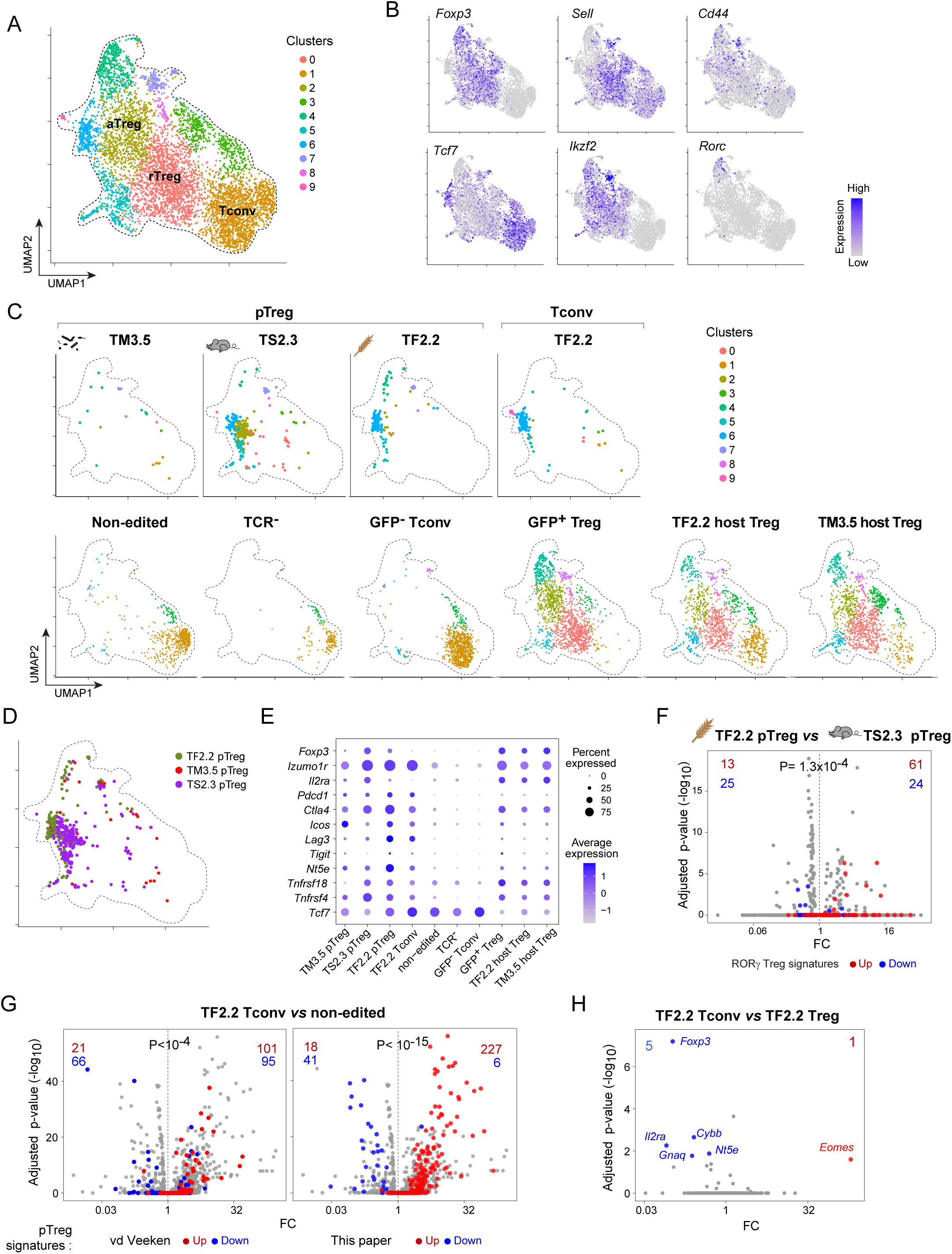
scRNA-seq reveals distinct transcriptomes in pTregs guided by different TCRs. Cells isolated from mLNs 10 days after adoptive transfer were analyzed by scRNA-seq analysis. Three TCRs were used: TM3.5 (EcN-reactive), TS2.3 (self-reactive) and TF2.2 (food-reactive) **(A)** UMAP of scRNA-seq data encompassing all groups, annotation based on expression of key indicator genes **(B)** and on reference cells in C. **(C)** Position of cells from each group on the collective UMAP; experimental groups on the top row, reference groups on the bottom row, including, from left to right: donor-derived non-edited and TCR-negative Tconv, standard colonic Tconv and Treg CD4⁺ cells from a Foxp3-GFP reporter mouse, and host Treg cells from TF2.2 and TM3.5 transfers (these contain a few Tconv cells because sorted on CD25). **(D)** UMAP plot grouping TF2.2-, TM3.5– and TS2.3-expressing pTregs. **(E)** Expression of Treg-related marker genes. The dot size represents the percentage of cells in a group that express the transcript, and the color indicates the average expression level. **(F)** Volcano plot of differentially expressed genes between TF2.2 and TS2.3 TCR-expressing pTregs. Red and blue dots represent genes highly expressed in colonic RORγ⁺ or Helios⁺ Tregs respectively. **(G)** Volcano plot of DEGs between TF2.2 TCR-expressing Tconv and non-edited donor cells. Red and blue dots: signature genes up-regulated in pTregs (left, from ref^13^, right, this paper) are highlighted in red, while genes downregulated in pTregs are shown in blue. **(H)** Volcano plot of differential gene expression between TF2.2 Tconv vs TF2.2 pTreg. Statistical significance of the enrichment in F and G was determined using Fisher’s test.

**Figs. 4C** also revealed that the donor-derived TF2.2⁺ cells that did not turn on FoxP3 (sorted as FoxP3-GFP-negative) did not fall as expected in the Tconv space of the UMAP, but instead very close to fully differentiated TF2.2 pTregs. They also expressed classic Treg markers like *Pdcd1*, *Ctla4*, *Lag3* or *Nte5* (**Fig. 4E**, **S5E**), but lacked the key *FoxP3* and *Il2ra* transcripts. Using an external signature of genes upregulated in pTreg cells in a lineage tracing system^13^, as well as a pTreg signature from our data (all donor-derived pTregs vs normal mLN Tregs from FoxP3-GFP reporter, **Table S4**), a comparison of TF2.2 Tconv relative to non-edited Tconv controls revealed that pTreg-differential genes aligned markedly with TF2.2 Tconvs (**Fig. 4G** **and S5F**), further supporting that those unconverted cells are highly Treg-like. Indeed, a direct comparison of transcriptomes of unconverted TF2.2 Tconv vs TF2.2 pTreg cells found essentially no difference, with the striking exception of *FoxP3* and *Il2ra* (**Fig. 4H** **and S5G**), and the upregulation of *Eomes*. We validated CD25 expression by flow cytometry and confirmed that it was restricted to FoxP3⁺ pTregs (**Fig. S5H**). It thus appears that these donor-derived Tconv that fail to convert actually encompass virtually all the changes leading to pTreg state, expect that they fail to induce *FoxP3* and *Il2ra*, almost reaching but not going beyond this tipping point.

### Do pTreg cells have memory ?

Once induced, how long do pTregs persist in the absence of the inducing antigen? In other words, does the pTreg pool contain a memory compartment, or is it continuously renewed? The questions speak to the nature of Treg memory^48–50^. It might be advantageous maintain memory Treg cells, providing rapidly recruited immunoregulation to prevent an over-exuberant secondary response by memory Tconv cells^51^, but it has also been proposed that transient “memory-less” Treg activity would avoid a cumulative state of generalized over-suppression^52^. Our experimental system allowed the ability to trace the fate of food-specific pTreg cells (**Fig. 5A**). After TCR editing and transfer of Tconv cells expressing either of two food-reactive TCRs into mice fed normal chow to elicit pTreg differentiation, part of the animals was switched to antigen-free amino-acid diet, and maintained for 6 to 10 weeks (we verified that, one week after such a switch, food antigen had cleared and was no longer able to elicit pTregs). With continued antigen presence, pTregs persisted in roughly the same numbers as at day10 (**Fig. 5B**). There was a clear drop of pTreg but not FoxP3⁻ Tconv population after the long period without antigen, which varied between tissues and with different TCRs: generally steeper for TF2.5 than for TF2.2, especially in the colon and SI LP for the former. Yet, in all instances, pTregs remained detectable in the absence of antigen, in timeframes commonly used for memory of conventional T cells. In terms of phenotype, TF2.2-driven pTregs kept their usual low/mid-level of RORγ⁺ proportions. On the other hand, TF2.5 pTregs, which achieve very high RORγ⁺ proportions, lost this almost completely in the SI and SI-draining mLN, while the phenotype was maintained in the colon and colon-draining mLNs, which we propose reflects the influence of high microbial load (**Fig. 5C**). These experiments clearly establish the existence of memory among food-reactive pTregs, and show that phenotypic changes can occur, independent of the initiating antigen.

**Figure 5.**
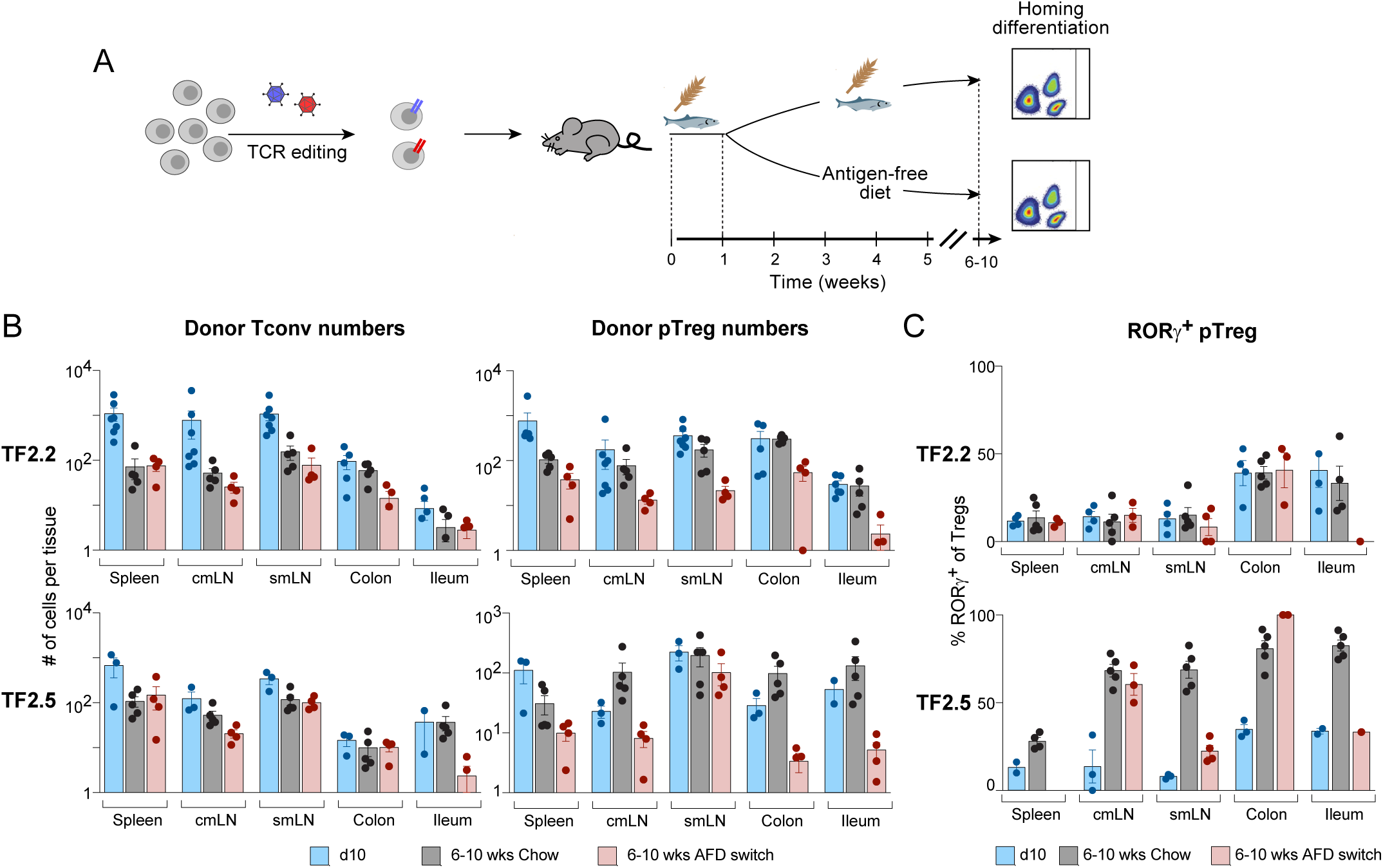
Memory in food-specific pTreg cells. (**A**) Experimental design. Tconv cells edited with food-reactive TCRs were transferred into a host fed normal chow for 1 week, then half the mice were switched to AFD for 6 or 10 weeks. **(B)** Quantification of cell numbers of TF2.2 or TF2.5 TCR-expressing Tconv or pTreg cells in indicated tissues. Day-10 data (blue bars) are from Figs. 2D/F. **(C)** Quantification of RORγ⁺ Treg frequencies in the same experiments. Each dot represents an individual mouse; bars represent mean, and error bars are standard error of the mean (SEM). *P < 0.05, **P < 0.01, and ***P < 0.001 (unpaired Student’s t test).

### Different antigen sources rely on different antigen presenting cells

Antigen presenting cells, and in particular dendritic cells (DC), exist in a spectrum types which differently influence the differentiation color of T cells to which they present antigens, including pTreg differentiation^53,54^. Recent studies using the Ovalbumin (OVA) tolerance model have shown the importance of RORγ⁺ APCs in regulating food-induced pTregs ^55–57^. In contrast, other studies reported that conventional DCs play a dominant role in OVA-responsive pTreg differentiation^58,59^. To address this contradiction, and to broaden the analysis beyond OVA-driven responses, we leveraged our pTreg-inducing TCRs to analyze the involvement of different APCs in pTreg differentiation in response to bacterial, food or self-antigens. In practice, we used as hosts two different lines of mice with selective antigen presentation defects. *Rorc^Cre^* x *H2-Ab1^fl/fl^*, hereafter *MHCII^△RORγ^*, lack MHC-II molecules in RORγ⁺ APCs, while *Clec9a*^cre^ x *H2-Ab1^fl/fl^* (*MHCII^△Clec9a^*) mice lack MHC-II expression in all cDC1 and most cDC2 cells ^55,60^

We transferred Tconv cell edited as above to express bacteria, food or self-reactive TCRs into these two strains (with cre-negative *H2-Ab1^fl/fl^* controls) mice (**Fig. 6A**). In the representative experiment of **Fig. 6B**, pTreg differentiation of TF2.2 expressing Tconv (food-reactive) was essentially abrogated in the *MHCII^△RORγ^* hosts, but only marginally affected in the *MHCII^△Clec9a^* host. Compiled data from several experiments (**Fig. 6C**) revealed very different outcomes according to the TCR/antigen combination. pTreg differentiation from both food-reactive T cells (TF2.2 and TF2.5) required MHC-II expression on RORγ⁺ APCs, but not on cDCs, while the outcome was exactly opposite for the self-reactive TS2.3 (no pTregs without cDC presentation, RORγ⁺ APCs irrelevant). Microbe-reactive T cells (TM3.5) were partially affected by both deficiencies. Thus, the involvement of different APC subsets in gut pTreg differentiation varies with the type of TCR/antigen pairs.

**Figure 6.**
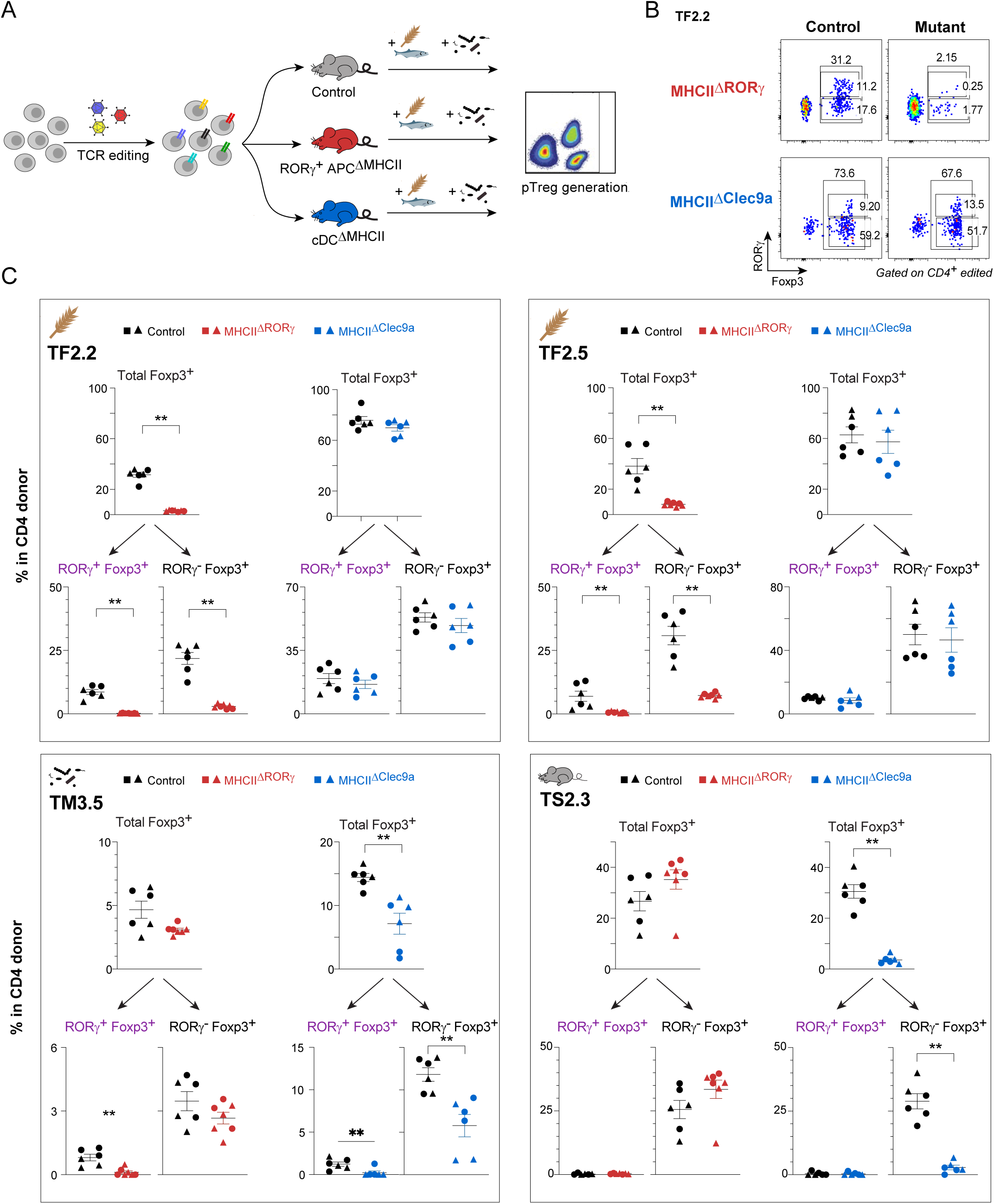
RORγ⁺ or cDCs APCs support pTreg driven by different TCR/antigen pairs. **(A)** Experimental design. TCR-edited Tconv cells specific for food, microbe or self-antigens, were transferred into hosts lacking MHC-II on RORγ⁺ APCs (red) or on cDCs (blue) or control littermates, and pTreg differentiation was assessed 10 days later. **(B)** Representative plots showing pTreg generation for TF2.2-edited cells in the indicated mutant host and control (gated on CD4⁺Edited⁺ cells. **(C)** Compiled quantification of mLN pTregs generated from four groups of TCR-edited Tconv cells, shown as whole FoxP3⁺ cells (as a proportion of donor cells, top panels), or split into RORγ⁺FoxP3⁺ and RORγ⁻FoxP3⁺ subsets. Each dot represents an individual mouse; bars denote mean ± SEM. Statistical significance determined by unpaired Student’s t test, **P < 0.01. Results are combined from two independent experiments.

The results were also interesting with regard to the phenotype of pTregs elicited in these partially-defective hosts. For food-reactive T cells in *MHCII^△RORγ^* mice, both RORγ⁺ and RORγ-negative pTreg were eliminated. The picture was more subtle for microbe-reactive TM3.5, the *MHCII^△RORγ^* deficiency preferentially affecting RORγ⁺ pTregs, the *MHCII^△Clec9a^* deficiency partially affecting both (and TS2.3 pTregs remained RORγ⁻ in either context) (**Fig. 6C**). These results indicate that it is not possible to make a 1-to-1 connection between the type of APC and the phenotype of pTreg whose differentiation it supports

## DISCUSSION

To explore how reactivity to different intestinal antigens controls T cell fate, more broadly than can be achieved with conventional transgenic mice, we performed a “mini-screen” by editing into CD4⁺ T cells a panel of TCRs originating from different types of colonic T cells, and with a range of antigen specificity. Transfer experiments with cells expressing these TCRs revealed a wide range in the ability to support pTreg differentiation and to yield RORγ⁺ vs Helios⁺ pTregs, depending on the type of cell from which the TCRs originated and the type of antigen they recognize. These results demonstrate the primacy of the TCR in determining CD4+ T cell fate at organismal barriers, and have implications for tolerogenic cell therapy.

The present results bring two important pieces of information concerning pTreg cells. First, pTreg cells are not necessarily RORγ⁺, and vice versa. We have previously argued^9^ that this relationship is not as obligate as often presented in the literature, but the present data should help close the debate: pTregs in our transfer experiments adopted a range of RORγ⁺ / Helios⁺ phenotypes. Most biased were pTregs driven by self-antigens, which were uniformly RORγ-negative, bacteria antigens promoted a majority of RORγ⁺ Tregs, while food antigens enabled both. Indeed, even for pTregs that express the same TCR, the proportion of RORγ⁺ Tregs was tissue-dependent, higher in the colon than in other tissues, and evolved over time. Secondly, pTregs are not only confined to the GALT, but disseminate widely in systemic lymphoid organs like the spleen or subcutaneous lymph nodes. Indeed, in absolute numbers, the spleen is where most pTregs reside, and they persist there long after differentiation. This dispersion may reflect continuous circulation of the pTregs, or trafficking outside of the GALT of APCs loaded with intestinal antigens. More generally, it ties in with the idea that T cell activation in the intestine can have unexpected consequences far outside the gut ^47,61–64^.

In addition, the profiling data provide an intriguing perspective on the transcriptional changes that occur in newly generated pTreg cells in an otherwise unperturbed host. These generally matched previous reports ^13^, but also brought out interesting divergences in pTregs driven by different TCRs/antigen combinations. The most arresting observation might concern those cells that expressed the same tolerogenic TCR but did not become FoxP3-positive, as evidenced by both reporter expression and RNAseq. We speculate that these represent an intermediate state reaching but not going through the tipping point of acquiring FoxP3. These cells might be related to the “wannabe” Treg-like cells that exist in FoxP3-deficient mice or humans, and also have many Treg characteristics ^65–67^. In this respect, pTreg differentiation might then behave much like thymic tTreg differentiation, with its late acquisition of *FoxP3* and *Il2ra* expression in intermediates that differentiate independently of FoxP3 ^68^.

The breadth of the TCRs studied here revealed striking differences between individual TCRs in directing local T cell fate. This observation differed from the notion that activation of any T cell by its cognate antigen in the gut could lead to pTreg differentiation, driven by unique conditions (high TGFβ, retinoic acid, etc) and APCs found there. TCRs originally isolated from Treg cells promoted pTreg development (although to variable extents), while Tconv-restricted TCRs largely failed to do so, cells maintaining a Tconv phenotype (with high proliferation in one case). Furthermore, TCRs derived from RORγ⁺ Tregs preferentially guided the differentiation of RORγ⁺ pTregs. There is a degree of circular reasoning in this notion that individual TCRs drive T cell differentiation back to their cell of origin, but it indicates that TCRs carry an intrinsic potential towards specific cell fates. A recent study of CD8⁺ T cells similarly showed that distinct clonotypes exhibit biased differentiation toward memory precursors or terminal effector cells, regardless of shared inflammatory cues or stochastic influences^69^. This influence likely reflects the TCR determining the antigen, the type of APCs that present the antigens, and the context in which activation occurs, as well as strength and kinetics of downstream signaling upon antigen recognition. However, the final outcome is also influenced by environmental cues. IL-6, previously found to modulate the proportion of RORγ⁺ Tregs ^6,15,45^, was required here for effective generation of RORγ⁺ pTregs.

Self-reactive Treg are generally believed to be generated in thymus. Our TCR editing allowed expression of self-reactive TCRs, all of which stemmed originally from RORγ-negative Treg cells, in Tconv cells, thus artificially bypassing normal thymic selection. We do not know if the cells from which these TCRs originated were thymically differentiated tTregs, or pTregs differentiated in the gut. Our results do establish, however, that peripheral tissue antigens that may have incompletely tolerized T cells in the thymus still have a chance of eliciting specific pTreg cells.

The APCs that drive the differentiation of pTreg and their subsets is an area of keen interest^31,55,57,58,70,71^. Our study highlighted the notion that different APCs are required for the generation of pTregs in response to different sources of antigen. At one end of the spectrum, differentiation of TS2.3 pTregs driven by self-antigen was wholly dependent on cDCs, with no requirement for RORγ⁺ APCs. Diametrically opposite, the impairment of RORγ⁺ APCs disrupted the generation of food-antigen-specific pTregs, affecting both RORγ⁺ and RORγ⁻ Treg subsets. This connection is consistent with and broadens several reports in the OT-II/oral-OVA model ^55,57,70,71^. Finally, pTregs induced with the TM3.5 TCR in response to *E. coli* antigen were intermediate, with a partial dependence on both cDCs and RORγ⁺ APCs. Previous studies had also shown a dominance of RORγ⁺ APCs in pTreg differentiation in the HH7-2/*Helicobacter* system ^31^. Our data suggest that, depending on the nature of the microbe, presentation of microbial antigens relies on different APC subsets to induce pTregs, in line with the earlier observation in *Listeria* vs *SFB* comparisons where T-cell effector functions were determined by the type of bacteria that produced the antigen, not by the antigen itself^72^.

In conclusion, the notion that a TCR drives the differentiation of uncommitted T cells towards the phenotype of the cell in which it was originally discovered has implication for T cell identity. It is not simply stochastically determined by encounters with cytokines, costimulatory molecules, or other non-specific influences, but to some extent pre-determined by the TCR itself. This notion has direct implications for T cell engineering: it does not suffice to equip a naïve T cell with a receptor whose specificity will drive it to a given location, but the use of a Treg-derived TCR will facilitate finding the proper antigen and APC to achieve the desired tolerogenic state.

## ACKNOWLEDGEMENTS

We thank Drs S. Ramanan and M. Sassone-Corsi who discovered the TCRs used here, Drs D. Kasper, E. Martens and D. Littman for provision of bacterial strains, K. Hattori, C. Araneo and I. Magill for help with mice, cell sorting and single-cell profiling. Dr. J. Chouinard and J. Espinal (LabDiet) for samples of diet components. This work was funded by NIH grants AI182126, AI150686 to CB&DM, from the Pew Trust, Parker Institute for Cancer Immunotherapy, CRISPR Cures for Cancer and Grand Multiple Myeloma Translational Initiative to JE, and an NIAID DP2 award (DP2AI171116) and a Pew Biomedical Scholar Award to CCB. VC was supported by a St. Baldrick’s Fellowship, WAN by the Swedish Society for Medical Research and the Swedish Research Council, CHW by the NSF Graduate Research Fellowship Program at UCSF, DLO was supported by a NIDDK T32 Postdoctoral Training Grant (T32DK007477) and a CRI Irvington Postdoctoral Fellowship (CRI4983).

## AUTHOR CONTRIBUTIONS

XC, CHW, YFP, AG, YC, VC, YC, JA performed the experiments; WAN, DLO provided materials and discussed interpretations; XC, CHW, WAN, YC, VC, CCB, JE, DM and CB designed the study, analyzed and interpreted the data; XC and CB wrote the first draft and all authors edited the manuscript.

## COMPETING INTERESTS STATEMENT

The authors declare no competing interests.

### Supplementary Materials

**Figs. S1 to S5**

**Tables S1 to S5**

## MATERIALS AND METHODS

### Mice

*Foxp^IRES-GFP^*/B6 ^73^, C57BL/6J (JAX: 000664), congenic Thy1.1 strain (JAX: 000406), Rosa26-*Cas9* knock-in mice (JAX: 026179), *Clec9a^cre^*, *Rorc^cre^*, *H2-Ab1^fl/fl^* ^60^ mice were maintained in our colony at HMS or MSK. HH7-2tg ^17^ transgenic mice were obtained from the Littman lab, BθOM^8^ mice were reconstituted from sperm from the MMRRC repository. All mice are under B6 background. Mice were housed under specific pathogen-free conditions. Gnotobiotic C57BL/6J mice were maintained at the Harvard Medical School Gnotobiotic Core Facility. SPF mice were under 5058 chow (LabDiet), gnotobiotic mice were fed on 5010 chow (LabDiet), or AFD (LabDiet, A12450KR) where indicated. The ingredients of each diet are in **Table S5**. All experimentation was performed following HMS IACUC protocol IS00001257, and MSK (IACUC protocol 21-05-007). Both male and female were used for all experiments. 7-12-week-old adult mice were used for most experiments, except *MHCII^△RORγ^* and *MHCII^△Clec9a^* hosts.

### TCRs

TCR used in this study were identified in colonic Treg and Tconv CD4⁺ cells from SPF or monocolonized mice, as described in Ramanan et al^29^, as detailed in figure S1.

### TCR editing of primary CD4⁺ T cells

TCR editing was performed largely according to Nyberg et al^36^. TCRs of interest were cloned as bi-cistronic cassette into the Ark313 vector, high-titer AAV stocks were prepared and used to infect CD4⁺ T cells from Cas9-exprssing transgenic mice, resulting in inactivation of both endogenous TCR loci and insertion of the recombinant TCR into the *Trac* locus for proper expression in T cells.

*TCR cloning into AAV vectors*: To generate TCR-targeting constructs for site-specific integration at the *Trac* locus, plasmids were derived by cloning an expression cassette for each TCR into pAAV-U6/gTrac-Trac-OT-I ^36^. designed to contain 500bp homology arms targeting *Trac* exon 1 flanking the intended knock-in region, with a U6 promoter-driven sgRNA targeting *Trac* for cleavage (UAUGGAUUCCAAGAGCAAUG) at the 5’ end, and a U6 promoter-driven sgRNA targeting *Trbc* (CUGGUGGGUGAAUGGCAAGG) for knockout at the 3’ end. The transgene cassette contained a P2A linker 1, TCRβ VDJ region coding sequences (starting from the first methionine codon of the V region gene), directly followed by *Trbc* sequences, P2A linker 2, TCRα VJ region coding sequences (starting from the first methionine codon from variable region sequencing), and partial TCRα constant region beginning with amino acid sequence “IQNPEP” which, in combination with the right homology arm (RHA), restores the complete endogenous TRAC coding sequence while forming a functional TCR of the intended specificity. Both *Trac* and *Trbc* sequences were codon-optimized to prevent re-cutting by their respective sgRNAs and eliminate sequence homology with endogenous genes to prevent recombination. Two distinct P2A ribosomal skipping sequences were used to prevent recombination between linker regions during cloning and integration. Individual TCRs were synthesized as double-stranded DNA and assembled using Gibson assembly and restriction enzyme cloning, with the AAV backbone generated exclusively through restriction enzyme cloning using BmgBI and PflMI sites to preserve AAV inverted terminal repeat integrity and avoid PCR-induced mutations that could compromise AAV packaging efficiency.

*Viral preparations*: AAV were produced using HEK293T cells. HEK293T cells were seeded in 150 mm dishes and transfected at approximately 70–80% confluency using polyethylenimine (PEI; Polysciences #23966). Each dish was transfected with 6 μg of AAV cargo plasmid containing the transgene cassette flanked by AAV2-ITRs, 8 μg of Ark313 capsid plasmid, and 11 μg of adenoviral helper plasmid providing essential adenoviral helper functions for AAV production. Transfection was performed in 200 μL of PEI per plate, and cells were incubated at 37°C with 5% CO₂. After 72 hours, viral particles were harvested from both the cell lysate and supernatant.

Cells were collected in AAV lysis buffer (50 mM Tris, 150 mM NaCl) and subjected to three rounds of freeze-thaw cycles to lyse the cells and release viral particles. Lysates were treated with Benzonase (25 U/mL; Millipore Sigma #70-664-3) at 37°C for 1 hour to degrade residual nucleic acids. Crude lysates were clarified by centrifugation at 3,000 × g for 10 minutes at 4°C, and viral particles were purified using iodixanol (OptiPrep, STEMCELL Technologies #07820) gradient ultracentrifugation. Following centrifugation, the AAV-containing fraction was extracted, dialyzed against PBS to remove iodixanol, and concentrated using Amicon Ultra-15 centrifugal filter units (Millipore Sigma #UFC910024).

AAV titers were determined by qPCR using DNase I (NEB #B0303S)-treated and Proteinase K (Qiagen #1114886)-digested AAV samples. qPCR was performed with SsoFast Eva Green Supermix (Bio-Rad #1725201) on a StepOnePlus Real-Time PCR System (Applied Biosystems #4376600) using primers targeting the viral genome. Relative viral genome quantities were estimated by comparison to a standard curve generated from serial dilutions of a plasmid of known concentration.

*Editing of primary CD4⁺ cells*: T cells were isolated from the spleens of Cas9-expressing transgenic mice. Spleens were mechanically disrupted and passed through a 40 μm cell strainer to obtain a single-cell suspension. Red blood cells were lysed using ammonium-chloride-potassium (ACK) lysis buffer, followed by washing with PBS. CD4⁺ T cells were enriched using the Dynabeads Untouched Mouse CD4 Cells Kit (Invitrogen #11416D), according to the manufacturer’s protocol. Purified T cells were activated with Dynabeads Mouse T-Expander CD3/CD28 (Gibco #11452D) and 200 U/mL recombinant human IL-2 (PeproTech #200-02) for 1 day. Activated T cells were resuspended at a density of 2 × 10⁶ cells/mL and incubated with AAV. The following day, the AAV-containing medium was replaced with fresh culture medium, and cells were maintained under standard culture conditions. Two days later, cells were resuspended in resting culture medium with 10 ng/mL IL-7 (BioLegend # 577802). Four days later, transduction efficiency and FoxP3 expression was assessed by flow cytometry before they were transferred into host mice.

### In vitro screening of antigen reactivity

*Antigen sources* included:

1. Heat killed bacteria. *E. coli Nissle* or *C. ramosum* were resuspended in PBS at the concentration of OD600=1, and then were heat treated at 75 °C for 1 hour. 5 μL of this suspension was used for each co-culture in a total culture volume of 150 μL.
2. Gut contents. Colon and cecum contents from indicated mice were diluted in PBS, vortex mixed, homogenized, filtered through 70 μm mesh, adjusted to concentration of OD600=1, and heat treated at 75 °C for 1 hour to kill all microbes. 5 μL of this suspension was used for each co-culture in a total culture volume of 150 μL.
3. Protein extracts. Whole 5058 chow or individual chow components (LabDiet) were dissociated in extraction media (600 mM KCl, 20 mM Tris-Cl pH 7.8, 10% Glycerol, and 1x cOmplete protease inhibitor in ddH2O). Colon contents were similarly processed and bacteria were removed by spin down at 2000g for 10 mins to extract digested food antigens. All samples then underwent 5 freeze thaw cycles alternating between dry ice and 37°C water bath. Samples were centrifuged for at 2000g for 10 minutes at 4°C. Supernatant was diluted into prechilled (–20°C) acetone at a 1:4 ratio, mixed thoroughly and incubated at –20°C for 30-60 minutes. Samples were centrifuged at 15,000g for 10min at 4°C. Supernatant was discarded and the remaining protein pellet was resuspended in DMEM without phenol red with 20 mM HEPES. Resuspended samples were centrifuged at 50g for 5 min at 4°C to remove any insoluble components. Protein concentration of the soluble fraction was measured by BCA assay. 20 μg/mL protein extracts were used for co-cultures.

*T-APC coculture*: To isolate APCs, spleens from B6 mice were digested with 0.5 mg/mL collagenase D and 0.1 mg/mL DNase for 30 min at 37℃. Single-cell suspensions were prepared, and CD11c⁺ cells were enriched by surface staining of CD11c-biotin antibody followed by Streptavidin MicroBeads (Miltenyi Biotec #130-048-101) treatment. Alternatively, whole adherent spleen cells were used as APCs. Single-cell spleen suspensions were plated onto 60 mm culture dishes for 90 min to allow APC to adhere. Non-adherent cells were removed by washing several times with complete medium. Fresh media were added for another hour, and adherent cells were then eluted by gentle pipetting. Similar results were gotten and combined from both methods. Enriched APCs were pre-activated with 100 ng/mL LPS with indicated antigen sources (20 μg/mL protein extracts or 5 μL heat killed bacteria or gut content in 150 μL total culture volume) overnight. Then APCs were co-cultured with TCR edited T cells at 1:1 ratio and co-responding antigens for 1.5 day. CD25 or Nur77 expression on TCRb⁺ cells were analyzed to indicated cell activation.

### Gene inactivation by electroporation of Cas9/gRNA complexes

CRISPR–Cas9 ribonucleoproteins (crRNPs) were prepared by incubating Cas9 protein with crRNA-tracrRNA duplexes, a pre-complexed two part-system with gRNA functionality. Five different predesigned crRNAs (CGTTGACAAGGGGGTTCTTC; ATGGATGACGCATTGGTACT; CTGTGCGTTGCAAACAGTGT; GGGGCAAATCAGGGTAACGG; CCCCACCAGGTGATCATTCA) targeting exon 2, 3, 4 and 6 of *Il6ra* were selected from Custom Alt-R CRISPR-CAS9 guide RNA service (IDT), and the common scaffold trans-activating RNA (tracrRNA) were obtained from IDT (Cat# 1072533). Non-targeting control crRNAs (IDT, Cat # 1072544, 1072545) were used as negative controls. The crRNA and tracrRNA oligos were mixed well in equimolar concentration to reach a final duplex concentration of 40 μM, and incubated at 37 °C for 30 mins. The crRNA-tracrRNA duplex were then mixed with Cas9-NLS (Macrolab, UC Berkeley) to achieve a duplex/Cas9 protein ratio of 2:1. For example, for 100 μL electroporation reaction, 11.25 μg Cas9 weas incubated with 0.4 nanomole of crRNA-tracrRNA duplexes. The complexes were allowed to form for 15 mins at 37 °C, and kept on ice before use (less than 30 minutes).

CD4⁺ T cells were enriched from Thy1.2-Cas9 or Thy1.1-Cas9 (CD45.2) mice by Dynabeads Untouched Mouse CD4 Cell Kit (Invitrogen, Cat# 11415D) per manufacturer’s instructions. 12 million primary CD4⁺ T cells were washed with Ca/Mg free PBS and centrifuged at 500g for 5 mins. The crRNPs complex were mixed gently with 100 μL electroporation solution cP4 (Lonza, V4XP-4024), incubated at room temperature for 2 mins, and cells were resuspended in this solution before transfer into Large Lonza cuvette strip. Nucleofection was then performed using Amaxa 4D-Nucleofector under X-P4 and program DS137. After electroporation, 750 μL of pre-warmed T cell culture medium supplemented with 10 ng/ml mIL7 were gently added to the cuvette containing electroporated cells. Electroporated cells were transferred into 12-well plates after 50 mins resting in 37 °C, 5% CO2 incubator, and cells underwent TCR editing 24 hours later.

### Bacteria culture and colonization

*Escherichia coli Nissle 1917* (Kasper lab collection) was grown overnight in Lennox Luria-Bertani (LB; BD Difco) supplemented with 100 μg/mL streptomycin at 37 °C. On the following day, EcN cultures were spin down 2000xg for 5 mins and resuspended in PBS to an OD600=1. To colonize SPF mice with EcN, adult SPF mice were gavaged with 20 mg streptomycin at day –1 and day 4 relative to cell transfer, and 100 μL EcN resuspension was gavaged on day 0 and day 5. Germ free C57BL/6 host mice were inoculated with 100 μL EcN suspensions, and maintained in Iso Positive Individually Ventilated cages. Two weeks after colonization, mice were used for cell transfer. For experiments using *MHCII^△RORγ^* and *MHCII^△Clec9a^* hosts, neonate mice were gavaged with EcN twice between at d10 and d15, and were transferred cells between d18-d26.

*Clostridium ramosum* (Kasper lab collection) was cultured on Brucella Blood Agar with Hemin and Vitamin K (Hardy Diagnostics #A30) under anaerobic conditions (80% N_2_, 10% H_2_, 10% CO_2_) at 37°C in an anaerobic chamber for one day. All colonies from one plate were resuspended in 1.5 mL PBS for oral gavage. Germ free C57BL/6 host mice were inoculated with 100 μL *Crm* suspensions, and maintained in Iso Positive Individually Ventilated cages. Two weeks after colonization, mice were used for cell transfer.

*Helicobacter hepaticus* was a kind gift from the Littman lab. It was thawed from Brucella broth and glycerol aliquots, resuspended in Brucella broth and cultured on TSA+5% sheep blood for 3 days anaerobically in an Embrient hypoxia chamber filled with O2-less gas (Airgas). It was then picked from the plate using cotton swabs and resuspended in Brucella broth+20% glycerol aliquots (OD600= >2) that were frozen in liquid nitrogen and thawed for gavage (200 μL).

*Bacteroides thetaiotamicron* was obtained from the Martens lab (U. of Michigan). It is grown on Brucella H+K agar for between 2 and 5 days. Multiple colonies are harvested and suspended in TYG medium, spun down and resuspended in TYG at OD600=1 to 1.5 before gavage (200 μL).

### Cell transfers and donor cell assessment

Unless otherwise specified, 100K-200K TCR edited Thy1.1/Cas9 cells were transferred into B6 mice via intravenous injection. In most experiments, tissues were analyzed ten days post cell transfer, except for Figure1E-F (one day) and Figure5 (6-10 weeks). The analyzed tissues included spleen, subcutaneous lymph nodes (scLN), small intestine– and colon-draining mesenteric LNs (smLN and cmLN), colon and ileum lamina propria. Donor cells were identified as live/dead-CD4⁺Thy1.1⁺. Among them, TCR⁻ cells were negative for TCRβ expression. TCR edited cells were positive for specific antibodies targeting the introduced TCR variable region, while non-edited cells were TCRβ⁺ but negative for edited TCR variable regions. For transgenic cell transfers, HH7-2tg and BθOM Tconv cells were sorted from the spleen as TCRβ⁺,CD4⁺ FoxP3-GFP⁻ and 10^5^ cells were transferred by intravenous injection.

### Lymphocytes isolation from tissues

Colon and ileum (removed Peyer’s Patches) were cleaned and treated with RPMI containing 1 mM DTT, 20 mM EDTA and 2% FBS at 37°C for 20 min to remove epithelial layers, minced and dissociated in collagenase solution in RPMI containing 1.5 mg/mL collagenase II (Gibco # 17101015), 0.5 mg/mL dispase (Gibco #17105041) and 2% FBS, with constantly stirring at 37°C for 40 min. Single cell suspensions were then filtered through 40 μm strainers and washed with RPMI solution. Lymph nodes and spleens were mechanically disrupted and filtered through 40 μm cell strainer to get single-cell suspensions. Red blood cells in the spleen were lysed with ACK lysing buffer (Gibco #A10492-01).

### Flow cytometry

Single-cell suspensions were stained with surface markers live/dead, CD4, TCRβ, CD25, indicated antibodies for variable regions of TCRα or TCRβ at 4°C for 20 min. For intracellular staining, cells were fixed in Fix/Perm buffer (eBioscience) at room temperature for 1 hour, followed by permeabilization in permeabilization buffer at room temperature for 1 hour in the presence of antibodies FoxP3, Helios, and RORγ. Cells were recorded by a Symphony flow cytometer (BD Biosciences) and analysis was performed with FlowJo software.

### scRNAseq pTreg profiling

*Sample preparation and sequencing.* scRNA-seq experiments were performed and analyzed as previously described ^74^. TCR edited cells (*Cas9/FoxP3^GFP^*) were transferred into CD45.1 hosts to analyze donor derived pTreg and Tconv cells. T cells infected with AAV carrying an early stopped codon in the TCR gene were used for TCR negative control (no TCR expression was confirmed). Cells subjected to mock infection (same in vitro culture procedure without virus) served as non-edited controls. For each TCR editing group, cells from 5-7 mice were pooled together to get sufficient cell numbers. Host cells from the same recipient mice, or FoxP3⁺ and FoxP3-cells from an extra *FoxP3^GFP^* mice were used for references.

Ten days post cell transfer, CD4⁺ T cells were enriched from mLN single cell suspensions before surface staining. Cells were then stained with antibodies against CD4, TCRβ, CD45.2, CD25, TCRVβ5 (TF2.2), TCRVβ12 (TM3.5), TCRVβ11 (TS2.3) and 4′,6-diamidino-2-phenylindole (DAPI) as a viability dye, along with hashtag antibodies (Biolegend, TotalSeq-C anti-mouse hashtags) for each population in staining buffer (DMEM, 5% fetal calf serum) for 20 min at 4°C in the dark. Indicated DAPI-CD4⁺ populations (CD45.2⁺TCRVβ5/11/12⁺GFP⁺ for donor pTregs, CD45.2⁺TCRVβ5⁺GFP-for TF2.2 Tconv, CD45.2⁺TCRβ-for TCR-, CD45.2⁺TCRβ⁺ for non-edited; CD45.2-GFP⁺ or CD25hi for host Treg cells, CD45.2-GFP-or CD25low for host Tconv cells) were sorted with a FACSAria III and pooled in a single collection tube at 4°C. All samples were pooled together, centrifuged, and resuspended in 2% Fetal Calf Serum (FCS).

Encapsulation was done on the 10X Chromium Controller (10X Genomics). Libraries were prepared using Chromium Single Cell 5′ Reagents Kit v2 according to the manufacturer’s protocol (CG000330). After cDNA amplification, the smaller fragments containing the TotalSeq-C–derived cDNA were purified and saved for feature barcode library construction. The larger fragments containing transcript-derived cDNA were saved for gene expression library construction. Libraries were sequenced together on the Illumina Novaseq X.

*Data analysis.* scRNA-seq data were processed using the standard CellRanger pipeline (10X Genomics), where RNA and Hashtag oligo (HTO) counts were obtained together through the cellranger count function. Data were analyzed in R using the Seurat package ^75^. HTOs were assigned to cells using the HTODemux function, and cells assigned doublets or negatives were eliminated from analysis. Cells with fewer than 1000 unique molecular identifiers (UMIs) or 400 genes and more than 2500 UMIs, 10,000 genes, and 5% of reads mapped to mitochondrial genes were also excluded from the analysis. Dimensionality reduction, visualization, and clustering analysis were performed in Seurat using the NormalizeData, ScaleData, FindVariableGenes, RunPCA, FindNeighbors (dims=1:30), RunUMAP (dims=1:30), and FindClusters functions. Cluster identity was determined based on expression of key marker genes (Fig. 4B and S5B). Differentially expressed genes (DEGs) were obtained using the FindMarkers function, with the cutoff based on absolute avg_log2FC>0.6, adjusted P value <0.05, and a pct>10%.

### Statistics

Data were shown as mean ± standard error of the mean. Statistical significance was determined using one sample t-test for Fig.1F, two-tailed t test (paired or unpaired as indicated) for two groups, and a fisher test for determining gene signature significance in volcano plots. Statistical significance was defined as P < 0.05. The level of significance in all graphs is represented as follows: * for P < 0.05, ** for P < 0.01, *** for P < 0.001. Statistics were performed in GraphPad Prism or R.

### Data availability

The scRNAseq data reported in this paper have been deposited in the Gene Expression Omnibus (GEO) database under accession GSE301231.

## SUPPLEMENTARY FIGURE LEGENDS

**Fig. S1.**
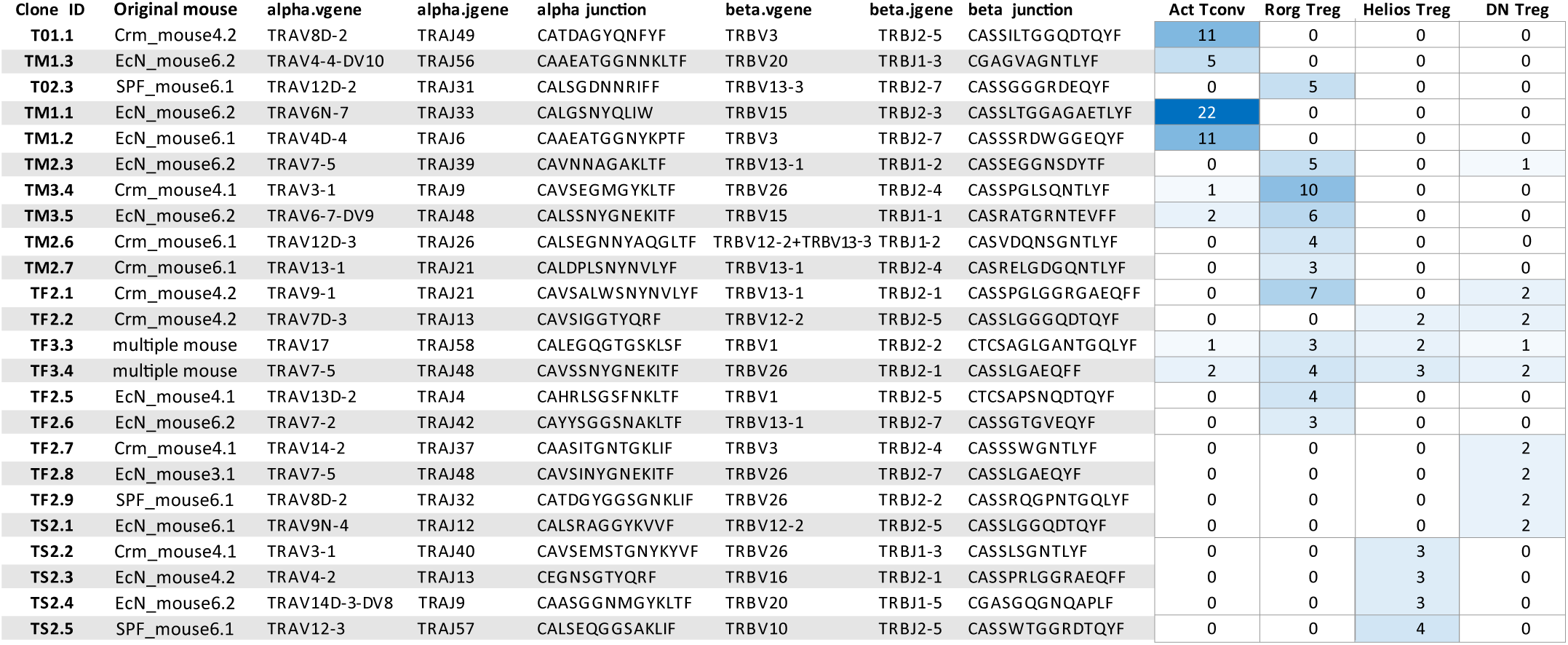
Summary table of selected TCR information, related to Figure 1. Amplified clonotypes were picked from Tconv, RORγ⁺Treg, Helios⁺Treg or RORγ⁻Helios⁻ (DN) Treg populations from gnotobiotic mice monocolonized by *C. ramosum* or *E.coli Nissle* ^29^. Numbers indicate the number of times each clonotype was recovered.

**Fig. S2.**
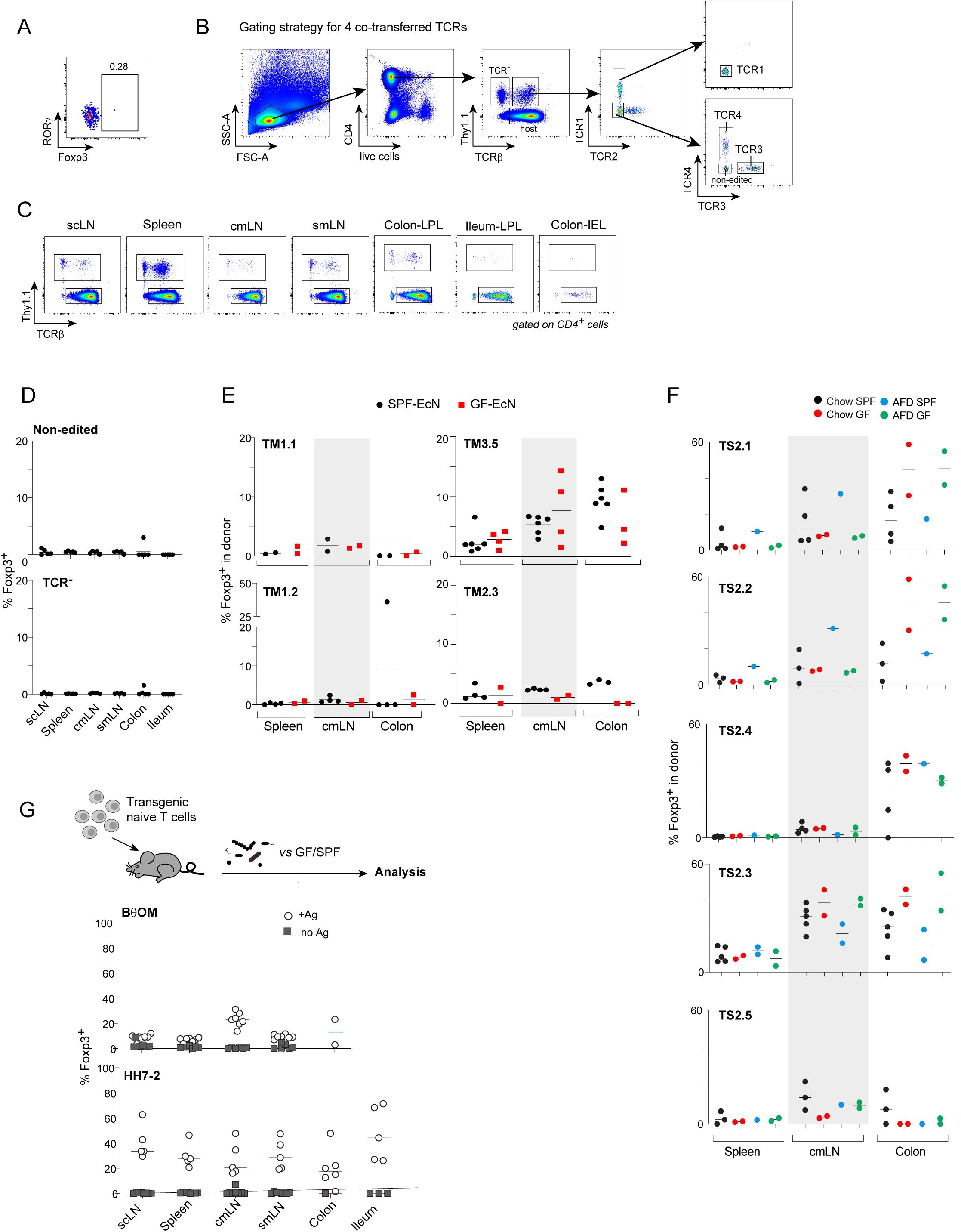
pTreg generation is consistent in different conditions, related to Figure 2. **(A)** Representative FACS plot of FoxP3 expression in TCR-edited cells before adoptive transfer. **(B-F)** CD4⁺ donor cells were transferred into indicated hosts for ten days. **(B)** Representative flow cytometric plots showing gating strategy to identify TCR-negative, TCR-edited and non-edited donor cells. **(C)** Representative flow cytometric plots of donor cell distribution in different tissues. **(D)** summary of FoxP3 expression in TCR-negative and non-edited cells. **(E)** Quantification of FoxP3 expression of TM3.5 and TM2.3 TCR-edited cells in EcN colonized gnotobiotic or SPF hosts. **(F)** Quantification of FoxP3 expression of TS2.1-32.5 TCR-edited cells in SPF mice under chow or AFD, or GF mice under chow or AFD diet. **(G)** Upper: Schematic illustrating experimental procedures for transferring naïve CD4⁺ TCR transgenic cells into hosts with or without corresponding bacteria colonization. Lower: Quantification of FoxP3 expression of BθOM or HH7-2 TCR transgenic CD4 T cells in B. theta or H. hepaticus colonized or control mice respectively. Each dot represents one mouse. Results are pooled from multiple independent experiments.

**Fig. S3.**
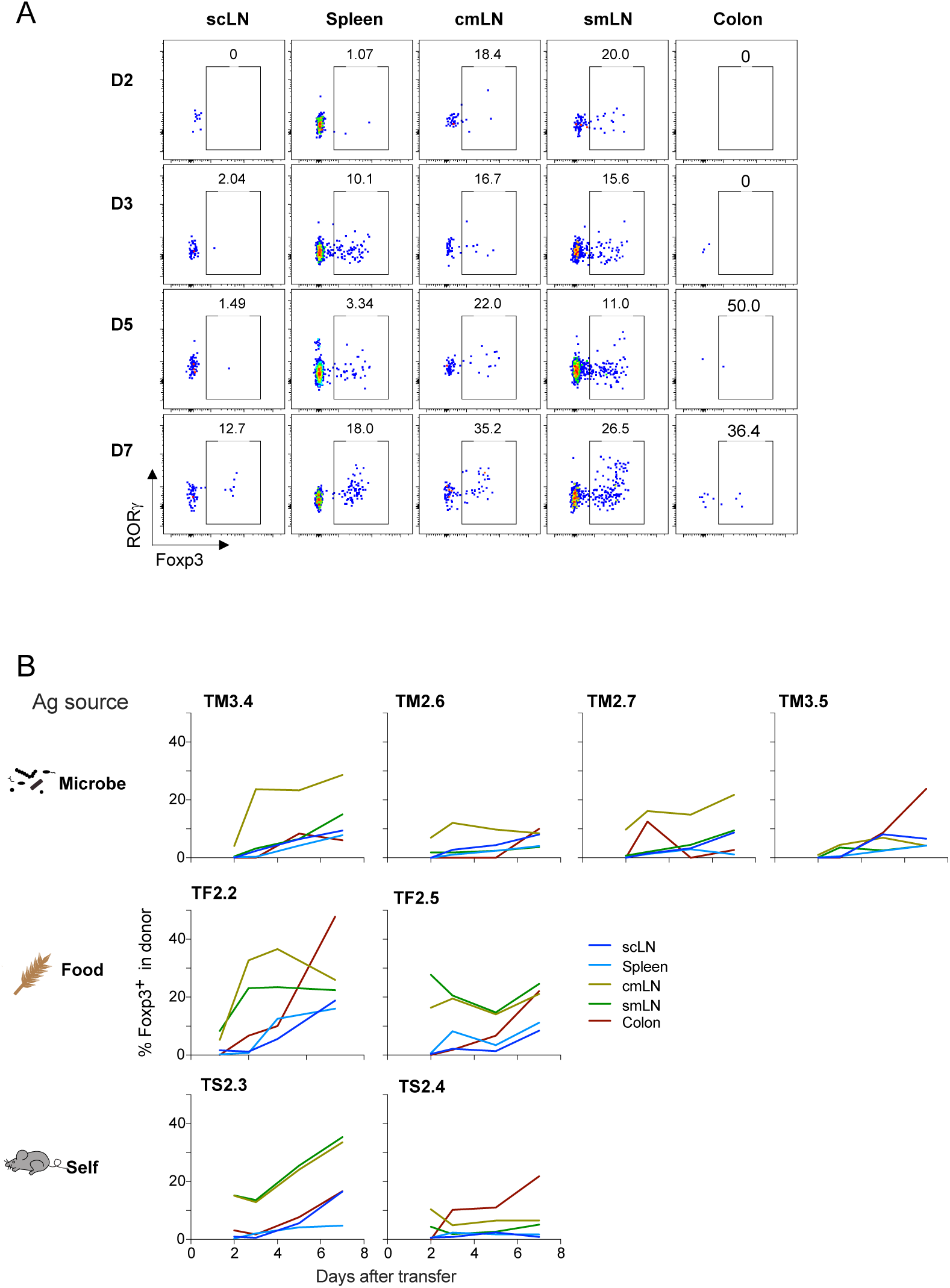
Time course of pTreg generation, related to Figure 2. TCR-edited CD4⁺ donor cells were transferred into B6 hosts and analyzed at indicated time point. **(A)** Representative FACS plots of FoxP3 and RORγ expression in TF2.5 TCR-edited cells. **(B)** Summary plots of FoxP3 expression. Results are pooled from two independent experiments.

**Fig. S4.**
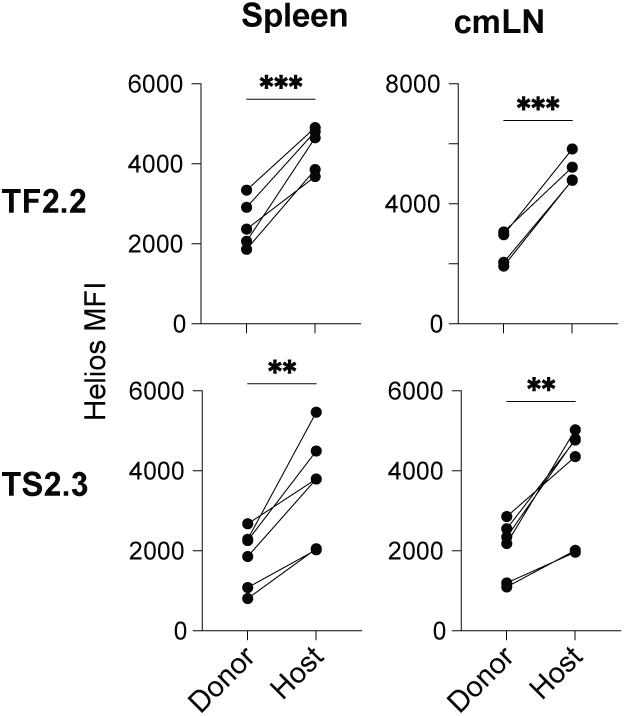
Helios expression levels in donor and host Tregs, related to Figure 3. TCR-edited CD4⁺ donor cells were transferred into B6 hosts and analyzed ten days later. Summary plots of Helios MFI in TF2.2– and TS2.3-pTregs and corresponding host Tregs. Results are pooled from multiple independent experiments. Each dot pair represent one mouse. Statistical significance determined by paired Student’s t test, **P < 0.01, ***P < 0.001.

**Fig. S5.**
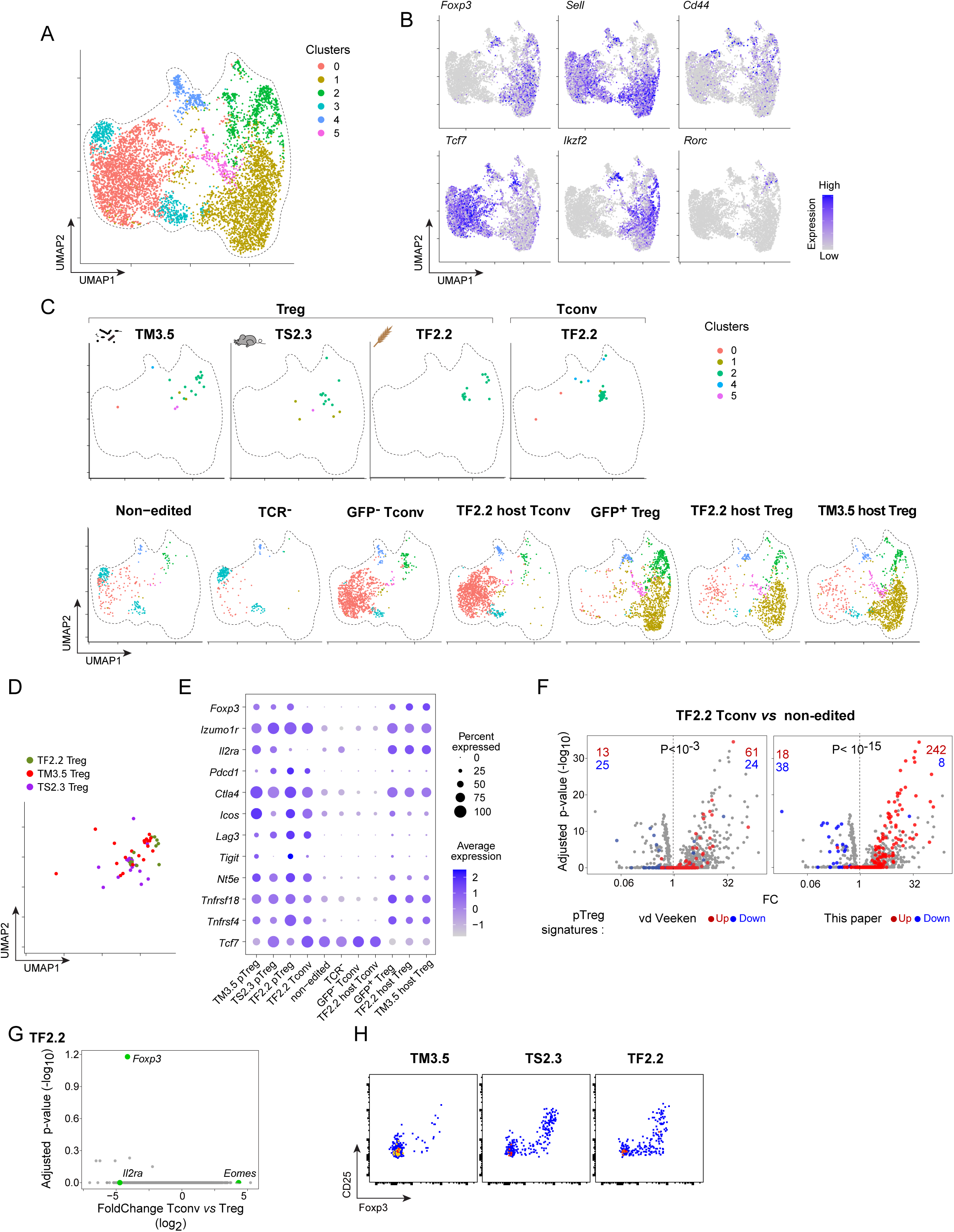
scRNA-seq reveals unique transcriptome of different TCR guided pTregs, related to Figure 4. scRNA-seq analysis of cells isolated from mLNs 10 days after adoptive transfer, including TM3.5, TS2.3 and TF2.2 TCR-expressing pTregs, TF2.2 Tconvs, non-edited donors, TCR-negative donors, FoxP3-GFP⁻ Tconvs, FoxP3-GFP⁺ Tregs and CD25 high Tregs from TF2.2 or TM3.5 reciepients. **(A)** UMAP of scRNA-seq data from all groups with semisupervised cluster annotation. **(B)** Feature plots showing key genes expression for Treg and Tconv annotation. **(C)** Individual UMAP plots of each group. **(D)** UMAP plot showing TF2.2, TM3.5 and TS2.3 TCR-expressing pTregs. **(E)** Dot plot of Treg related marker genes expression. The dot size represents the percentage of cells in a group that express a given feature, and the color indicates the average expression level. **(F)** Volcano plot of differential gene expression between TF2.2 TCR-expressing Tconv and non-edited donor cells. Genes upregulated in pTregs (based on reference dataset ^13^) are highlighted in red, while genes downregulated in pTregs are shown in blue. Red and blue numbers indicate the counts of pTreg upregulated and downregulated genes that are also differentially expressed in TF2.2 Tconv or non-edited cells, respectively. Statistical significance of the enrichment was determined using Fisher’s test. **(G)** Volcano plot of differential gene expression between TF2.2 Tconv vs TF2.2 pTreg. **(H)** Representative FACS plots of CD25 and FoxP3 expression in TM3.5, TS2.3 and TF2.2 TCR-expressing cells from mLN.

## SUPPLEMENTARY TABLES

**Table S1:** detailed CD25 MFI FC and Nur77 expression related to figure 1.

**Table S2:** detailed donor cell tissue distribution and pTreg generation related to figure 2 and 5.

**Table S3:** DEGs between TF2.2 and TS2.2 pTregs.

**Table S4:** pTreg up– and down-regulated gene list.

**Table S5:** Detailed diet information.

